# Panoramic visual statistics shape retina-wide organization of receptive fields

**DOI:** 10.1101/2022.01.11.475815

**Authors:** Divyansh Gupta, Wiktor Młynarski, Anton Sumser, Olga Symonova, Jan Svatoň, Maximilian Joesch

## Abstract

Statistics of natural scenes are not uniform - their structure varies dramatically from ground to sky. It remains unknown whether these non-uniformities are reflected in the large-scale organization of the early visual system and what benefits such adaptations would confer. Here, by relying on the efficient coding hypothesis, we predict that changes in the structure of receptive fields across visual space increase the efficiency of sensory coding. We show experimentally that, in agreement with our predictions, receptive fields of retinal ganglion cells change their shape along the dorsoventral retinal axis, with a marked surround asymmetry at the visual horizon. Our work demonstrates that, according to principles of efficient coding, the panoramic structure of natural scenes is exploited by the retina across space and cell-types.

## Introduction

The idea that sensory neurons exploit the statistical structure of natural stimuli to minimize the metabolic cost of information transmission has been a guiding principle in neuroscience for over half a century ^1–3^. This conceptual framework, known as the efficient coding hypothesis ^4^, has provided successful theoretical accounts of sensory coding across species and sensory systems ^5–8^ with the retina being the paramount example ^9^. Most of the work in the retina has been focused on retinal ganglion cells (RGCs), the neurons that relay visual information from the eye to the brain. It has been demonstrated that multiple properties of RGCs: the shape of receptive fields (RF) ^10–13^, organization of RF mosaics ^14,15^ and the ratio of ON to OFF RGC cell-types ^16^, can be explained as adaptations to the natural sensory environment. In all of the mentioned cases, *ab initio* theoretical predictions about efficient encoding of natural scenes have led to a better understanding of the physiological and anatomical properties of the retina.

One way a sensory neuron could implement an efficient code is by removing predictable (or redundant) components from sensory stimuli, in a transformation known as predictive coding. This prominent hypothesis suggests that the center-surround structure of RGC RFs is a manifestation of such design principle ^10^. According to this hypothesis, the surround computes a prediction of the stimulus value in the center of the RF. The predicted value is then “subtracted” from the center through inhibition, which dramatically reduces the amount of neural resources used to convey the stimulus downstream. Predictive coding and related information-theoretic principles ^11–13,17^ typically assume that the structure of natural scenes is uniform across the visual field. However, as demonstrated recently, local contrast and luminance vary prominently across the elevation within the natural visual field of a mouse ^18,19^. Such systematic variation affects the signal-to-noise ratio (SNR) of the input to RGCs. To understand how this inhomogeneous noise structure should shape RGC RFs, we developed a simple, predictive-coding model. When adapted to natural statistics of mouse vision, our model generates three key predictions linking the shape of optimal RFs and their position within the visual field. First, the relative surround strength should increase with increasing elevation, due to a consistent increase in brightness from the dim ground to the bright sky. Second, the center size should decrease along the same axis. Third, due to a rapid change of signal intensity between lower and upper fields of view, RFs centered on the horizon should have strongly asymmetric surrounds, with the upper half being stronger than the bottom one.

To test these predictions experimentally, we established a novel system that enables recording and characterization of the RF structure at high resolution, at the scale of thousands of RGCs in a single retina. Such technological development enabled us to collect a dataset of 31135 RGC RFs covering the entire central retina, which was crucial to test our theory. We found a close agreement between theoretically optimal RF architecture and the variation of RF shapes across the retina, suggesting that RGCs exploit global asymmetries of natural scenes for maximizing coding efficiency. Furthermore, we explored these adaptations across the diversity of functional RGCs types ^20^; each thought to share the same physiology, morphology, intra-retinal connectivity ^21–24^. We identified a systematic dorsoventral variation of the RF shape, regardless of the functional type. Finally, we show that these global adaptations are preserved in awake-behaving animals with intact eyes. Our results thus indicate that adaptations to the panoramic natural statistics structure retinal representations used by the brain.

### Efficient coding predicts organization of RF shapes across the visual field

To understand how the statistical structure of natural scenes shapes receptive fields (RFs) across the visual field, we developed a model of sensory coding in retinal ganglion cells (RGCs; Fig. 1a). Our approach is closely related to the predictive coding (PC) theory, which postulates that RGCs recode outputs of photoreceptor cells to minimize the metabolic cost of sensory information transmission ^10^. Following this theory, we modeled neural responses as a linear combination of the RF and natural stimuli, distorted by different sources of constant noise, e.g., biochemical or synaptic ^9,25,26^ (Fig. 1a, right panel). The computation performed by such RF can be understood as a difference of the weighted center of the stimulus and its surrounding neighborhood. Our model generates predictions consistent with PC (Supplementary Fig. 1) as well as related theories of efficient sensory coding ^11,13^.

**Fig. 1.**
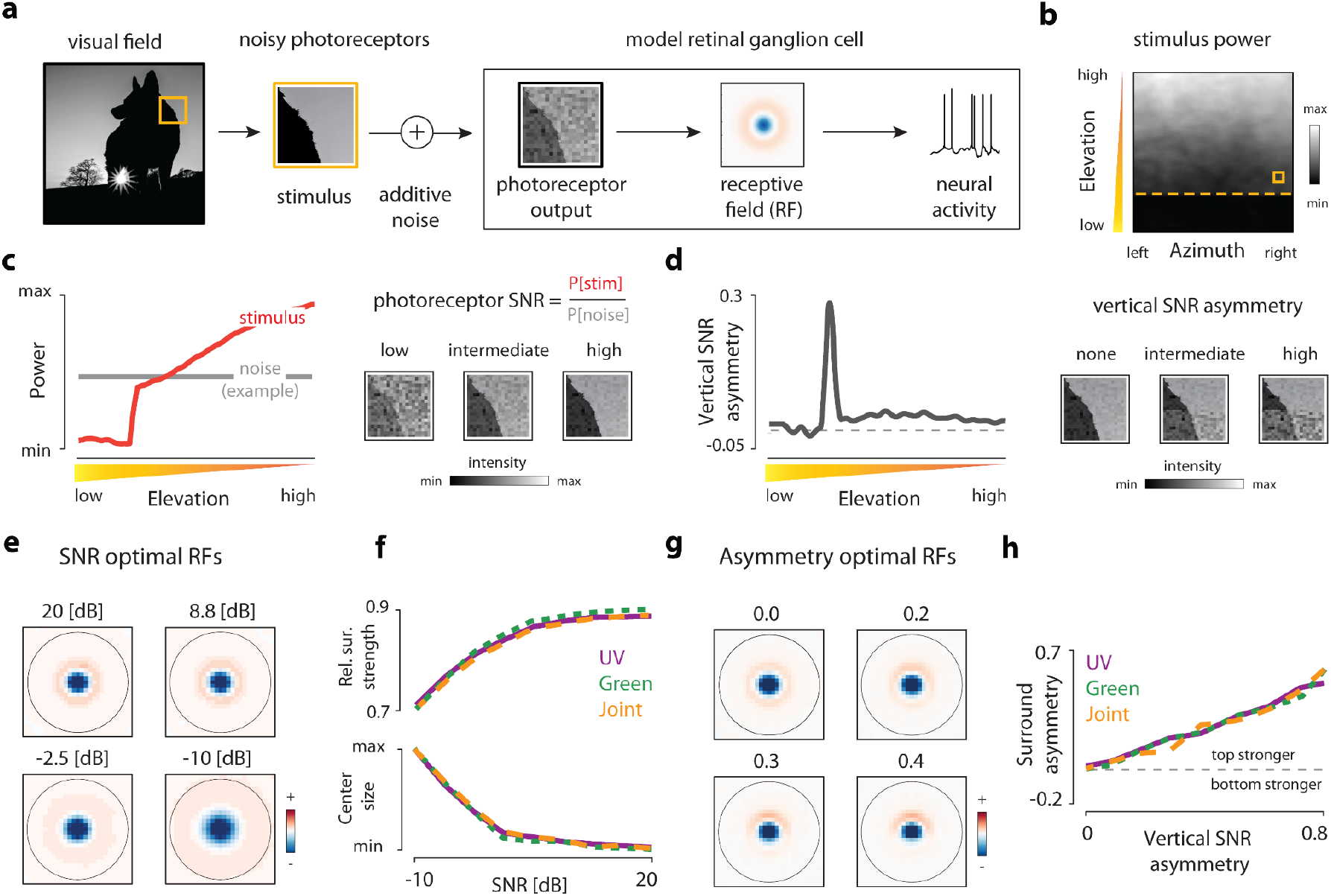
Predictive coding and natural scene statistics. (**a**) Schematic of the linear model of a receptive ganglion cell encoding noisy photoreceptor outputs. (**b**) Average stimulus power in the mouse field of view in the ultraviolet range (natural image data - courtesy of Hiroki Asari ^18^). Orange dashed line denotes the simulated horizon. Orange frame illustrates the size of the model receptive field. (**c**) Stimulus power in the ultraviolet range (left panel, red line) and example noise power level (left panel, gray line) as a function of elevation in the visual field. Increasing stimulus power increases the SNR (right panel, illustration). (**d**) Vertical SNR asymmetry in the ultraviolet range as a function of elevation in the visual field (left panel). Change in SNR asymmetry is due to asymmetric power in the stimulus at the horizon line (right panel - illustration). (**e**) Predictive coding RFs optimal for different levels of signal-to-noise ratio. Receptive fields were smoothed with a 2 × 2-pixel window for display purposes. (**f**) Relative surround strength (top panel) and center size (bottom panel) of optimal predictive coding RFs increase and decrease respectively, with increasing photoreceptor SNR. Purple, green and orange lines correspond to the UV, green, and joint spectrum respectively. (**g**) Predictive coding RFs optimal for different levels of vertical SNR asymmetry. Receptive fields were smoothed with a 2 × 2-pixel window for display purposes. (**h**) Surround asymmetry of optimal predictive coding RFs increases with increasing vertical SNR asymmetry of photoreceptor output. Line colors analogous to panel (**f**).

Predictive coding theory of the retina assumes that statistics of natural stimuli are stationary across the visual field ^10^. However natural scenes are spatially inhomogeneous. To understand this inhomogeneity, we examined a set of natural images collected specifically to study mouse vision ^18^. In agreement with previous studies ^18,19^, we found that the power of the light intensity decreases gradually with the elevation and drops-off suddenly close to the simulated horizon line (Fig. 1b,c). Under the assumption of constant noise level, the SNR of photoreceptor outputs (i.e., RGC inputs) should therefore follow the analogous pattern (Fig. 1c, gray line depicts example value of constant noise power). Moreover, due to an abrupt change of the stimulus power, stimuli centered at the horizon yield a highly non-uniform SNR pattern (Fig. 1d). To find how such inhomogeneities could affect sensory representations across the retina, we numerically optimized RFs to minimize the strength of neural responses averaged across a set of natural image stimuli (see Methods).

The shape of the optimal RF depends on the relative strength and the structure of noise. When the SNR decreases, the center of the optimal RF broadens, and the surround becomes more diffused (Fig. 1e), independent from the absolute SNR. This qualitative change is manifested in increasing relative surround strength (Fig. 1f, top panel) and decreasing center sizes (Fig. 1f, bottom panel). The optimal RF shape is additionally modulated by the spatial pattern of SNR (Fig. 1g). When the SNR is spatially non-uniform (e.g., when the signal is stronger in the upper half of the stimulus), the optimal RF becomes asymmetric (Fig. 1g). This effect is particularly visible in the increasing asymmetry of the surround as a function of SNR asymmetry (Fig. 1h). Because of such systematic variation of the stimulus power across the visual field, the PC model predicts three qualitative links between the position of a neuron in the retina and the shape of its RF. First, the strength of the RF surround relative to the center should be increasing with elevation across the visual field. Second, the size of the center should increase in the opposite direction. Third, RFs located at the horizon should have surrounds that are substantially stronger in the upper than in the lower half. Such distribution of RF shapes would indicate that RGCs exploit global statistics of the visual field to maximize the efficiency of sensory coding. These three predictions stand in contrast to the dominant view that RGC RFs are uniform across the retinal surface. Furthermore, predictions of the PC model are reproducible across different ranges of the light spectrum (Fig. 1f,h), sets of natural stimuli (Supplementary Fig. 1) and depend primarily on weak assumptions about the correlation structure of natural images ^10^ (Supplementary Note 1, Supplementary Fig. 2). We thus consider them to be a robust consequence of the efficient coding hypothesis.

### Large-scale characterization of receptive fields across the retina

Testing these theoretical predictions requires a high-resolution characterization of RGC RFs from extended regions of the retinal surface. Currently, however, it is not practical to perform such large-scale characterizations with any of the existing methods. Multiphoton imaging approaches can measure large numbers of RGCs^20,27^, but only at a moderate throughput (~150 RGCs at a time ^20,27^). Multi-electrode array recording approaches have improved this number but are limited by the recording area that is placed on top of the electrode array ^15^. Moreover, RF estimates generated by current approaches lack a clear surround structure ^20^. To circumvent these limitations, we designed a high-throughput and low-cost epi-fluorescent approach that enables imaging larger field-of-view (FOV) (1.7 mm^2^ sampling at ~1 μm/pixel) and permits >1 hr long recordings of the same FOV while avoiding artifacts caused by small retina wrinkles and laser scanning. Our method takes advantage of red Calcium sensors (e.g., RCamp1.07 ^28^) that separate the Ca^2+^ indicator’s red-shifted excitation light from the opsin absorption spectrum (Fig. 2a,b; see methods), and allows robust responses to UV visual stimulation. We used the VGluT2-cre driver line to specifically target RGCs (Fig. 2c), leading to a uniform expression across the entire retina. All RCamp1.07 positive somata correspond to RGCs, as seen by the RGC specific marker RBPMS ^29^ (Fig. 2d). Double-positive cells accounted for ~40 % of all RGCs. This expression pattern appears to be RGC-type-specific, as seen by the SMI-32 alpha-RGCs co-labeling. Alpha-RGCs were consistently excluded from the expression profile, apart from a single, sparse and spatially distributed type (Fig. 2e). Using this line, we were able to reproduce and expand previous large-scale imaging results in single retinas, as seen in direction-selective (DS) and non-DS responses (Fig. 2f), the cardinal DS response distributions (Fig. 2g), and clustering and reproducibility of responses to changes in frequency, contrast and luminance, known as the “chirp” stimulus ^20^ (Fig. 2h & Supplementary Fig. 3). By sequentially recording 3-7 FOVs (Fig. 2c, orange box depicts one FOV), each for approximately 25 min, we were able to record neural activity from up to ~6 mm^2^ of retinal surface (~40 % of the total retinal area). By experimental design, the position of each FOV was random. Moreover, the strength of the functional responses in each consecutive session was unaltered (Fig. 2i). Importantly, using a novel “shifting” white-noise approach, where the checker positions are randomly shifted to increase the RF spatial resolution (Supplementary Fig. 4 & see methods, and ^30^), we were able to estimate high-resolution and high-SNR spatiotemporal RFs for ~85 % of recorded cells (Fig. 2j, top row). The quality of these RF estimates allowed for automatic parameterization of the spatial RF into center and surround using a difference of Gaussians model (Fig. 2j, bottom row & Fig. 2k, Supplementary Fig. 5, see methods). In total, we recorded 11 retinas, reconstructing and parameterizing 31135 spatiotemporal RFs, enabling an unprecedented opportunity to index RGC responses across single retinas. This methodology will enable functional developmental screens and circuit dissections due to its simplicity, efficiency, and affordability, extending the current retinal research toolbox.

**Fig. 2.**
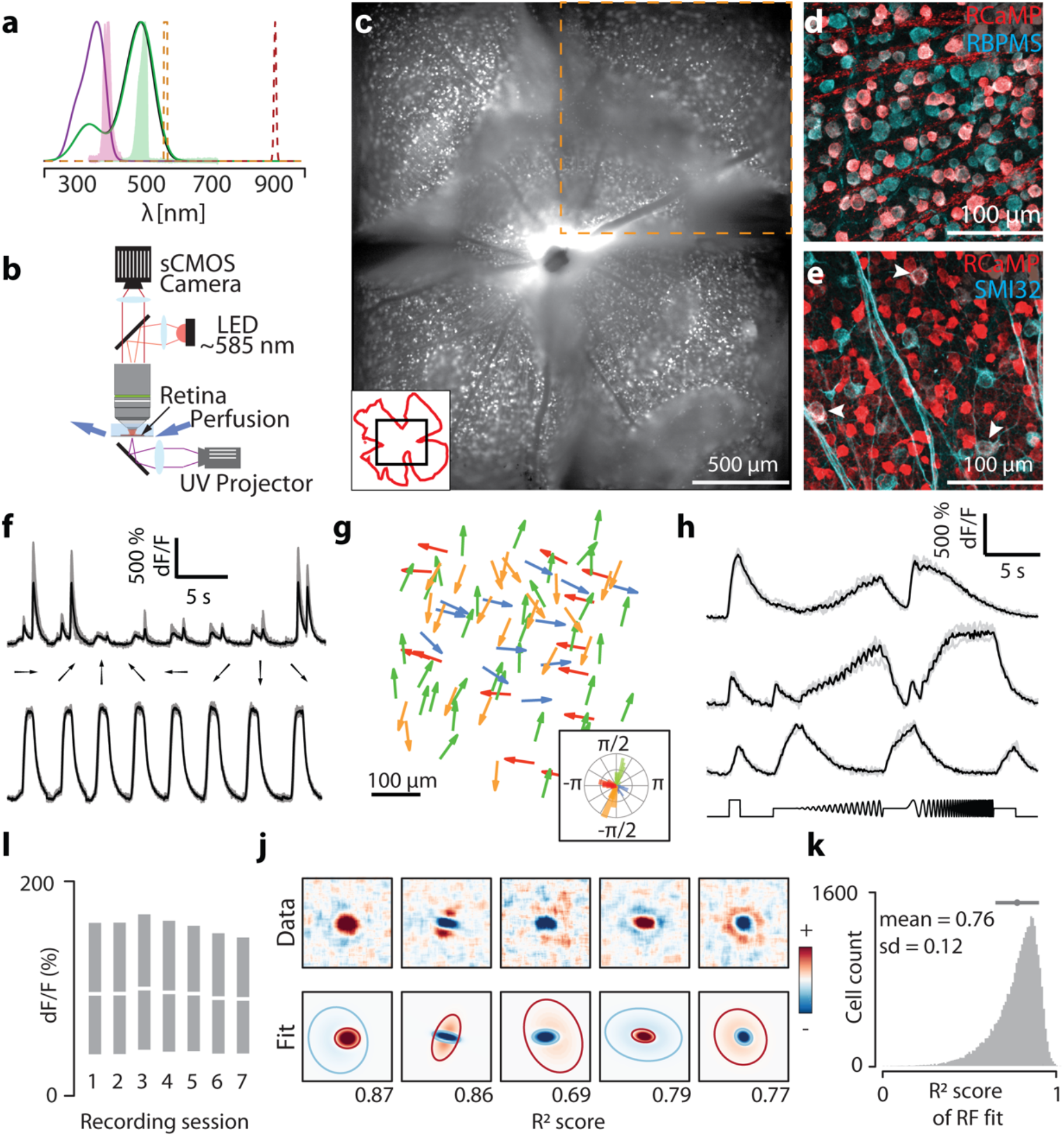
Large-Scale retinal receptive field mapping. (**a**) Normalized absorption spectra of mouse photoreceptors (purple: S opsin; green: M opsin). Overlaid, the normalized emission spectra of the UV and green light emitted by the DLP projector (filled purple: UV, filled green: green stimulus light), epi-fluorescent (orange) and two-photon (red) excitation. (**b**) Schematic of our epi-fluorescent imaging setup. (**c**) Montage of five consecutively recorded fields (orange inset denotes one field) of a whole-mounted mouse retina from a Vglut2-ires-cre; TITL-R-CaMP1.07-D; ROSA26-ZtTA triple transgenic mouse. Inset - black: imaged montage, red: retinal outline. (**d**) Double-label immunostaining of RCamp1.07 expressing RGCs (red) and RBPMS (cyan). (**e**) As in (**f**) but labeling SMI32 (cyan). Arrowheads depict double-labeled cells. (**f**) Example Ca^2+^ signals (grey: 5 repetitions; black: mean) from direction (top panel) and non-direction selective RGCs (bottom). (**g**) Example distribution of preferred directions in one FOV. Inset: polar plot of DS preference. (**h**) Example Ca^2+^ signals to chirp stimulus from 3 different RGCs (grey: 5 repetitions; black: mean). (**i**) Recording Ca^2+^ signal stability across sequentially imaged FOV for nine retinas (each session lasts ~25 min, 3-7 sessions per retina) White lines and gray bars denote medians and 25^th^ and 75^th^ percentile range of the dF/F distribution respectively. (**j**) Example receptive fields recorded using “shifting” white noise (top) and their respective parametrizations (bottom). Blue and red ellipses correspond to two standard-deviation contours of the ON and OFF Gaussians, respectively. (**k**). Histogram of goodness of fit for all recorded receptive fields.

### Receptive fields are adapted to anisotropic scene statistics across the visual field

Taking advantage of the high-resolution RFs, we examined variations in RF shapes and strengths across the retina. Given that the PC model does not determine the polarity of the optimal RF and to globally compare all retinal RFs, ON-center RFs were flipped in sign, such that all centers were negative and all surrounds were positive. This allowed us to pool across all cells within small bins on the retinal surface and visualize the average spatial RF at different locations of the retina (Fig. 3a, Supplementary Fig. 6). We observed a general and reproducible trend across 11 retinas: a streak-shaped area where all RF surrounds were oriented towards the optic nerve, and below which, hardly any RF surrounds were observed (Fig. 3a, Supplementary Fig. 6). To compare these spatial variations of RFs with our theoretical predictions, we oriented 6 of the recorded retinas to a common coordinate system using the immunohistochemically determined S-opsin gradient. Average RFs in the ventral, central-dorsal and peripheral-dorsal retina (Fig. 3b, bottom row) qualitatively matched model RFs predicted for the upper, medial and lower visual fields, respectively (Fig. 3b, top row). To confirm the change of relative surround strengths independently from surround asymmetry, we computed the radial profiles of RFs and these also strongly resembled the radial profiles for model RFs (Fig. 3c). Overall, model RFs qualitatively reproduced all aspects of average spatial RFs across different elevations with remarkable detail.

**Fig. 3.**
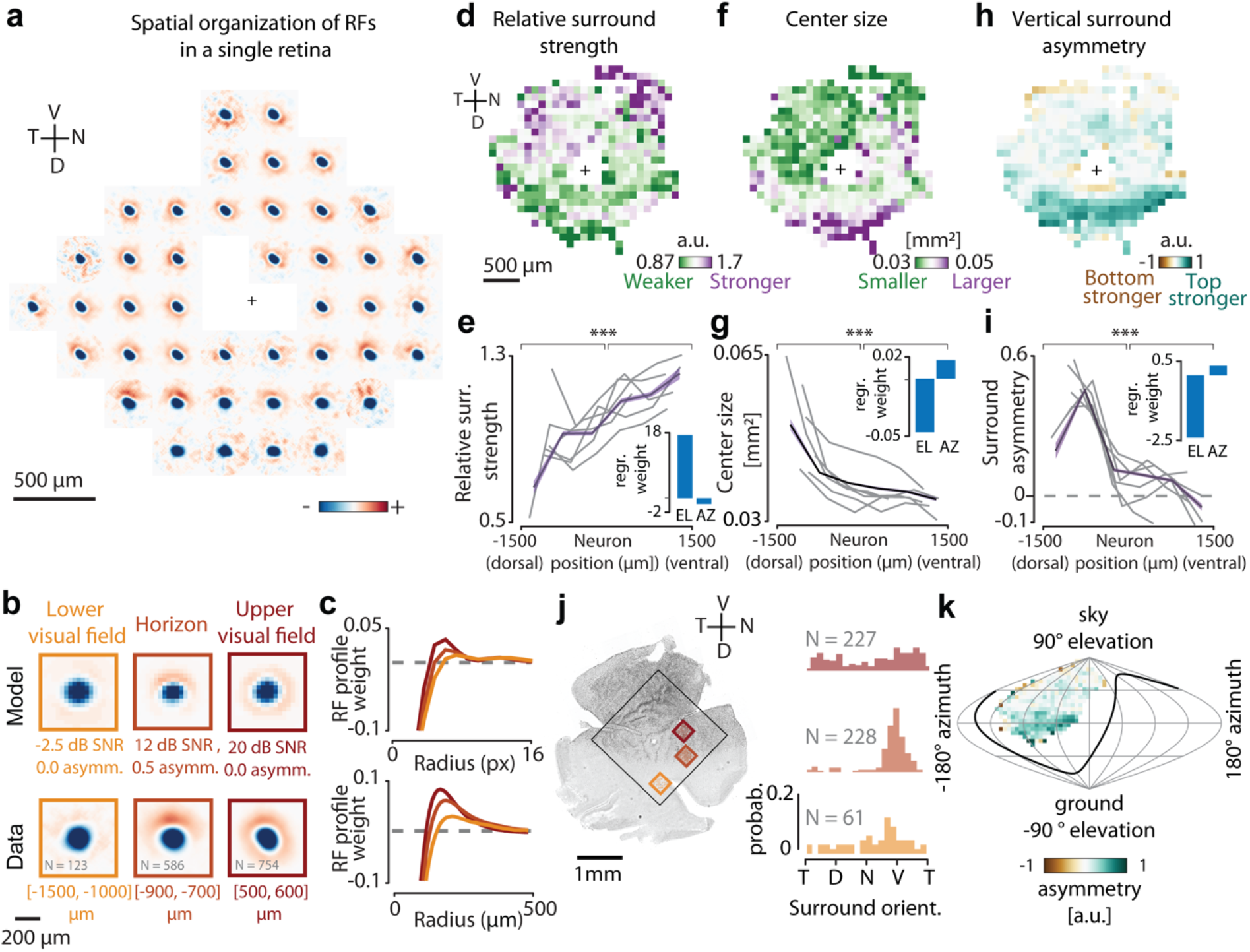
Retina-wide receptive field architecture. (**a**) Average spatial RFs of all RGCs pooled from square bins of size 300 μm at different positions of one retina (n = 64 ± 52 cells per bin, Black cross: optic nerve head). (**b**) Top row: Optimal RFs predicted by the model at different elevations of the visual scene. Bottom row: Average spatial RFs of neurons along different dorsoventral locations on the retina. (**c**) Top: Radial profiles of model RFs at different SNR levels. Bottom: Mean radial profiles of RGC RFs in bins along the dorsoventral axis. (**d**) Mean relative surround strengths of RGCs within 100 μm bins, pooled from n = 6 retinas. (**e**) Relative surround strengths for RGCs within 6 equally spaced bins along the dorsoventral axis (color: mean and SEM pooled from all n = 15686 RFs, grey lines: individual retinas, inset: linear regression weights of RF parameter on elevation (EL) and azimuth (AZ)). (**f** & **h**) Same as (d), but for center size and vertical surround asymmetry, respectively. (**g** & **i**) same as (e), but for center size and vertical surround asymmetry, respectively. (**j**) Left: One of the retinas, immunostained for S-Opsin. Black box shows the region imaged for RF mapping. Right: Normalized histograms of surround orientations of RGCs within corresponding bins marked on the left. (**k**) Data from (**h**) overlaid on a sinusoidal projection of visual space (n = 6 retinas). The animal is centered at (0 ° latitude; 0 ° longitude), facing towards the viewer and the black line shows the area of the visual field viewed by one eye. (p-values for Kolmogorov-Smirnov test: (**e**) 6.11×10^−5^, (**g**): 2.84×10^−4^, (**i**): 1.16×10^−5^; see Supplementary Fig. 7 for extensive statistical comparisons). V - Ventral, N - Nasal, D - Dorsal, T - Temporal.

To measure these phenomena quantitatively, we made use of the RF parametrizations and pooled RF parameters for all cells in different 2-d (Fig. 3d-h) or 1-d (Fig. 3e-i) bins across the retina. In line with our theoretical predictions (Fig. 1f, top panel), our analysis shows that the relative surround strength increases gradually along the dorsoventral axis (Fig. 3d), a trend visible in every single retina (Fig. 3e). Next, we explored if we could observe any global change in RF center size. As predicted (Fig. 1f, bottom panel), center sizes decrease only across the dorsoventral axis (Fig. 3f,g, Supplementary Fig. 7). While examining the spatial distribution of differences in upper and lower halves of the RF surrounds, we identified a consistent and prominent asymmetric streak in the dorsal retina, between 700-900 μm dorsally from the optic nerve (Fig. 3h,i), as one would expect from asymmetric visual inputs (Fig. 1e). Accordingly, linear regressions weights are substantially stronger in elevation, but not azimuth, for all three trends (Fig. 3e-i (insets), Supplementary Fig. 7). Overlaying the measured RF asymmetries with the opsin gradient indicated that the asymmetry is pronounced in the opsin transition zone (Fig. 3j, right panels). To test whether this streak corresponds to the horizon line within the animal’s visual field, we transformed the retinal coordinates to visual coordinates ^31^ and used the S-opsin gradient ^32^ to define the dorsoventral axis (Fig. 3j,k, see methods). In visual coordinates, the center of this asymmetric streak is located at 0 ° elevation, spanning the entire azimuth of our imaged FOVs (Fig. 3k), in line with our theoretical predictions (Fig. 1c). Finally, these trends also align in 5 additionally imaged retinas, where the true orientation could not be determined by the opsin gradient (Supplementary Fig. 8).

### Adaptations to natural scene statistics across retinal pathways

It has long been assumed that specific RGC pathways have stereotyped response properties, shaped by the interactions between direct excitation in their center and indirect inhibition in their surround ^33^. Thus, one would expect that these center-surround interactions are uniform across visual space for most RGCs pathways. To assess the specificity of the observed RF adaptations across functional RGC pathways, we clustered cells based on the temporal RF profile of 31135 RGCs into functional groups using a Gaussian Mixture Model (GMM), as done previously ^20^. Consistent with the proportion of RGCs labeled in our line (Fig. 2d), we defined 10 functional clusters (Fig. 4a). Chirp responses were not used for clustering, since we observed a strong bias for RGCs with weak surrounds (Supplementary Fig. 3c,d). Each cluster group had distinctly shaped temporal filters, corresponding to different functional properties such as ON or OFF selectivity, transient or sustained responses, and monophasic or biphasic selectivity, as seen in their average profiles (Fig. 4a, right panels). Cluster membership statistics were conserved across different retinas (Supplementary Fig. 9). Moreover, the relative positions of RGCs belonging to individual clusters tile the retina in a mosaic-like arrangement in many cases (Fig. 4b, left panels), confirming that some clusters are indeed functionally distinct and irreducible RGC types ^15^. As expected, many clusters represent a combination of RGC types that cannot be identified solely by their temporal profiles (Fig. 4b, cluster 10, see Supplementary table 1 for tiling statistics). We next used this classification and looked at the relative surround strength, center size, and asymmetric strength across clusters (Fig. 4c). As with the global pooling of RF shown in Fig. 3, cells in each functionally defined cluster increase their relative surround strengths gradually in the dorsoventral axis and decrease their center sizes accordingly. Moreover, all clusters contribute to the asymmetric streak (Supplementary Fig. 9c), consistent with the distribution of asymmetries in the opsin transition zone, where most cells have a strongly oriented surround with a ventral bias (Fig. 3k). All three trends were statistically significant for all clusters in elevation, but not azimuth (Supplementary table 1). These results indicate that a substantial proportion of RGC pathways adapt to the constraints imposed by natural statistics.

**Fig. 4.**
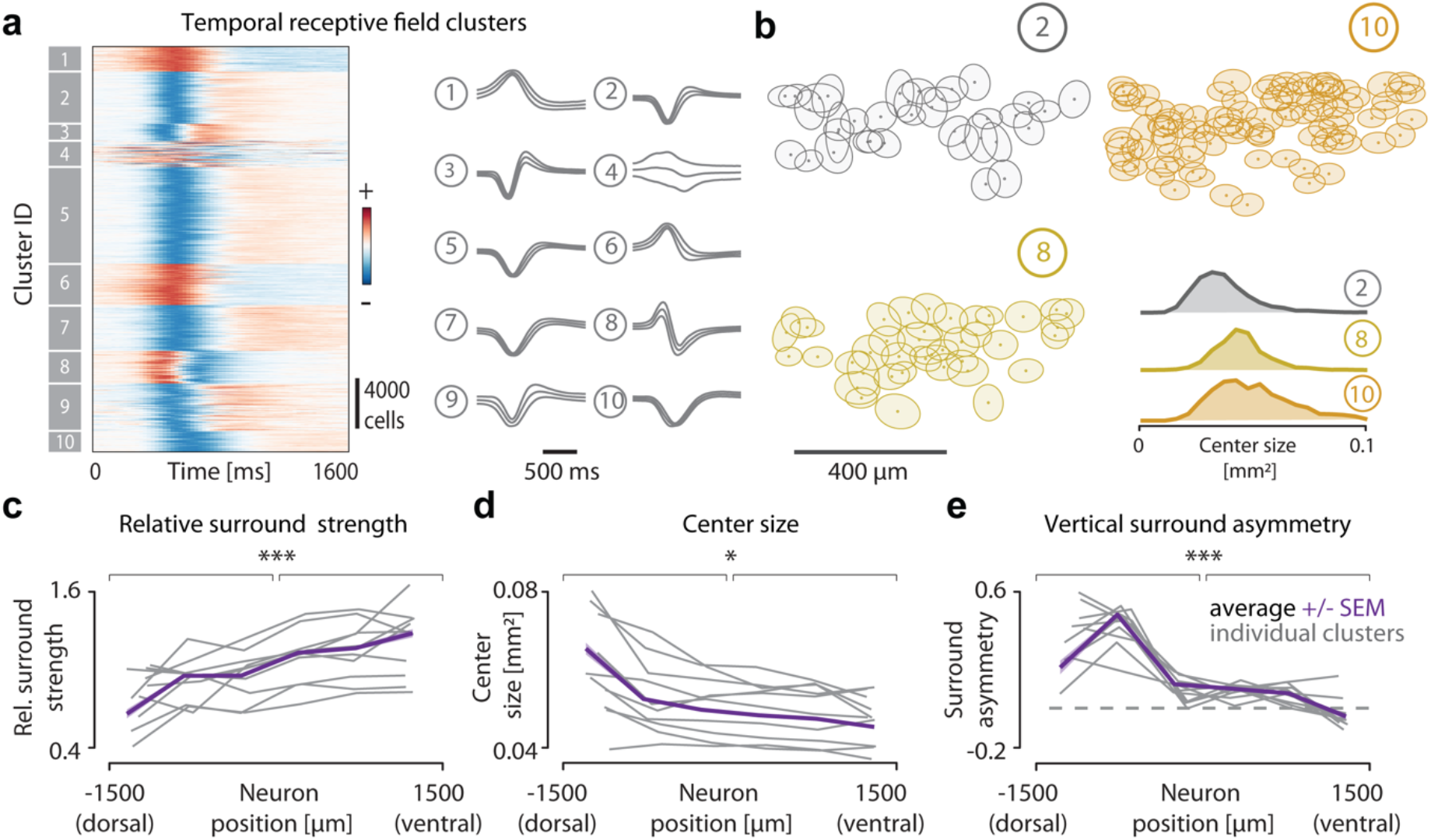
Global adaptation across retinal pathways. (**a**) Left: Temporal RFs of 31135 RGCs, grouped by cluster membership. Right: Mean and standard deviation of temporal RFs in each cluster. (**b**) Locations of RGCs (dots) and their RF centers (filled ellipses – 1 standard deviation of the center Gaussian) belonging to three different clusters in a small region of one retina. Bottom right: Distribution of center sizes for these three clusters. (**c**) Trends for relative surround strength (left), center size (middle) and vertical surround asymmetry (right) for cells within each cluster (grey) and pooled from all clusters (color), binned as in Fig. 3E. (p-values for Kolmogorov-Smirnov test: left: 6.17×10^−5^, middle: 0.02, right: 2.62×10^−7^).

### *In vivo* panoramic receptive field anisotropies match theoretical predictions

To test whether the adaptations to the panoramic visual scene statistics affect sensory coding during behavior in naturalistic conditions, we decided to map RFs of the RGC axonal terminals in the superior colliculus (SC) (Fig. 5a,b). These experiments have the advantage of testing our hypothesis in retinas that retain the complete adaptation machinery, from an attached pigment epithelium to functioning pupil constriction. Moreover, they provide important additional insights into the effects of the M-cone and rod pathways, which are saturated in our *ex vivo* retinal imaging system. For this purpose, we expressed the Calcium indicator GCaMP8m ^34^ in RGCs, using adeno-associated viruses (AAV) in 3 mice. Subsequent implantation of a cranial window above the SC allows for visualization of RGC axonal terminal activity with 2-photon Calcium imaging in awake, behaving mice. The field-of-view varied per recording from 0.32 to 1.85 (median = 0.68) mm^2^ of superficial SC surface (Fig. 5b), encompassing 23 to 57 (median = 41) visual degrees in elevation. GCaMP8m expression spread homogeneously across SC (Fig. 5c). Using the same previously used “shifting checker” stimulus, we recorded 53000 terminals in total, reconstructing 10000 RFs above our quality index (see methods). Compared to the RFs recorded from *ex vivo* retinas, *in vivo* measured RFs were blurred along the main axis of saccadic movements (Fig. 5d, 2-d RFs). Thus, to compare across animals, we aligned the RFs of each animal to their respective saccadic plane, which due to the head fixation had one consistent axis parallel to the ground to each animal as shown previously ^35^ (Supplementary Fig. 10) and corrected the mouse head position to match the retinal coordinates (see methods). To avoid any bias due to eye movements, we then used the 1-d profiles (1-d RFs) of the orthogonal axis for further analysis (Fig. 5e, 1-d RFs). By binning and averaging 1-d RFs along the latero-medial axis, spanning from lower to higher visual field (Fig. 5fg-i, Supplementary Fig. 11a), the three predicted trends became visible: (i) the surround strength increases, (ii) the center size reduces, and (iii) the surround becomes more symmetric. Similarly to our retinal results (Fig. 3b), the mean 2-d RFs qualitatively matched model RFs predicted for the upper, medial and lower visual fields, respectively (Fig. 5f). Next, we analyzed the RF parameters on mean 1-d RFs across visual space. As with previous results, the RF properties were significantly different above and below the horizon and have substantial regression weights on elevation (Fig. 5g-i, insets) but not on azimuth (Supplementary Fig. 11d-i). Consistently, the average visual maps (Supplementary Fig. 12) resembled the ones measured *ex vivo* (Fig. 3d-h). Thus, our *in vivo* results provide independent corroboration that the visual system is adapted to the constraints imposed by the panoramic natural statistics.

**Fig. 5.**
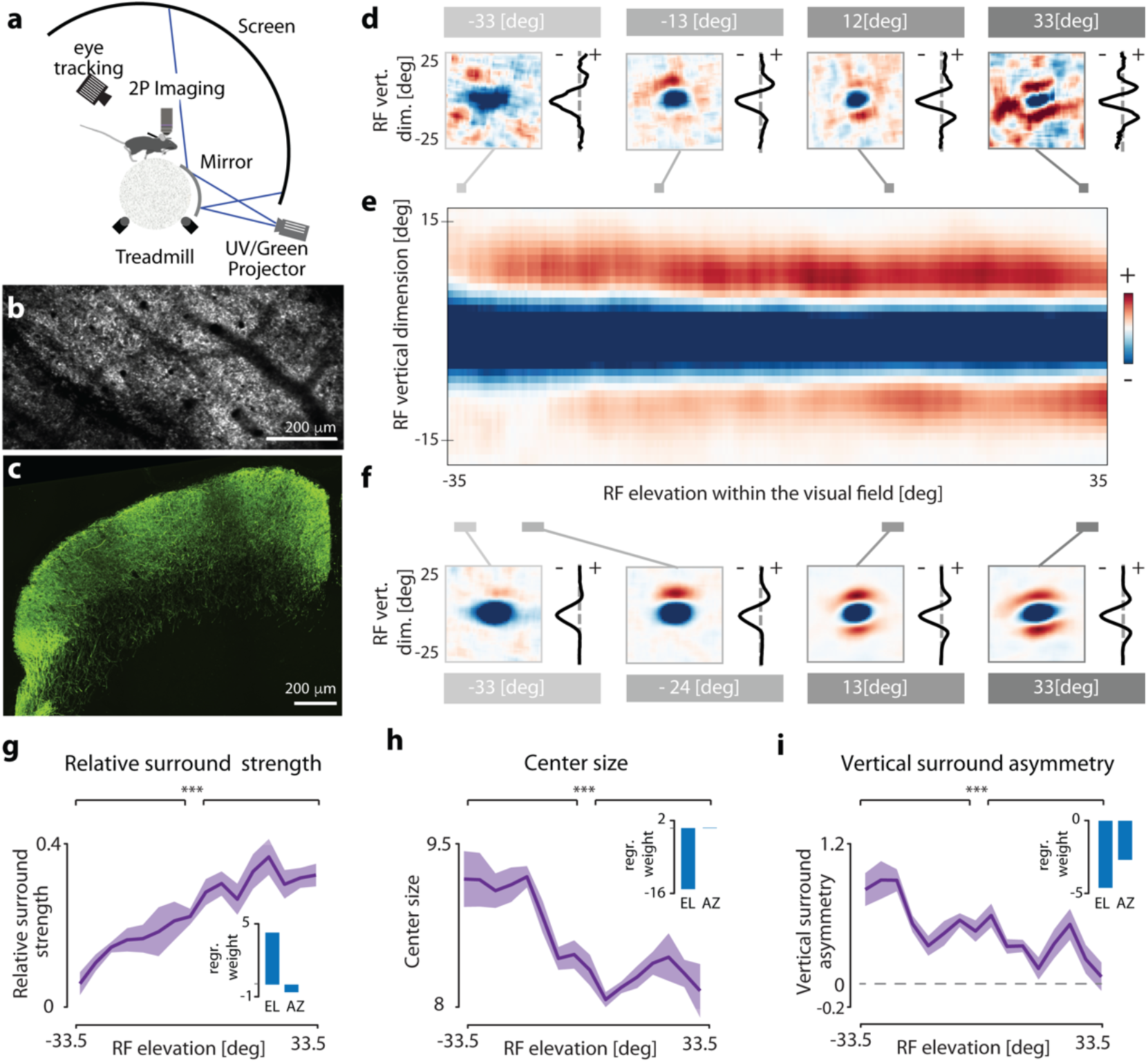
Colliculus-wide RGC’s receptive field architecture. (**a**). Schematic of *in vivo* multiphoton imaging setup. (**b**) Field of view of a standard multi-photon recording of RGC axons expressing GCaMP8m in the superior colliculus (max. projection). (**c**) Immunostaining of GCaMP8m of an example coronal section of the superior colliculus, showing homogenous RGC labeling across the visual layers (green). (**d**) Example RGC bouton receptive fields recorded using “shifting” white noise (left) and their respective vertical 1-d center profiles (1-d RFs, right) at different elevation levels (gray lines). (**e**) average 1-d RFs, in 0.22 ° bins over elevation (smoothed horizontally in 5° gaussian window for display purposes). (**f**) Example average receptive fields binned at 4.1 ° visual angle (left), with their respective 1-d RFs (right) at different elevation levels (gray bars). (**g**) Relative surround strength of 4.1 ° binned and parameterized average 1-d RFs, shading indicating SEM across azimuth bins (Supplementary Fig. 11a-c). Inset: linear regression weights of individual bouton (n = 9810) 1-d RF parameter on elevation (EL) and azimuth (AZ). (**h, i**) As (g), but for center size and vertical asymmetry, respectively. (p-values for Kolmogorov-Smirnov test: (g) 2.91×10^−10^, (h): 1.16×10^−12^, (i): 1.48×10^−06^).

## Discussion

In this study, we leveraged a novel, high-throughput neural imaging setup (Fig. 2) to identify a new kind of adaptation to panoramic scene statistics in the retina. In agreement with theoretical predictions derived from the efficient coding framework (Fig. 1), our experimental results indicate that the global RF architecture is adapted to encode panoramic natural scenes efficiently (Figs. 3 & 4). These results were further corroborated in RGC terminals of awake animals (Fig. 5), indicating that panoramic, efficient representations impact downstream processing during behavior. Classically, RGCs are known to dynamically change the strength of the RF surround in response to varying light levels ^36,37^, which is thought to further increase the efficiency of sensory coding ^9,38,39^. Our findings demonstrate that, in addition to such dynamic effects, RF shapes are also determined by static factors, namely their position within the visual field. In that way, the retina simultaneously exploits the large-scale spatial and fine-scale temporal structure of the visual space.

How could the visual system establish such global RF architecture without fine-tuning each RGC pathway independently? One partial mechanism would be the non-uniform distribution of spectral sensitivity across the retina ^32^. Such distribution has been discussed to be relevant for color vision ^40^, contrast coding ^19,41^, and the detection of aerial predators in the sky ^42^, but simultaneously, could influence the static RF adaptations. For example, whereas the mouse retina has green light-sensing photoreceptors across the entire retina (M- and Rod-opsin), UV sensitivity follows a sharp dorsoventral gradient (S-opsin) ^32^. From the RGC’s perspective, both inputs will be added at mesopic conditions, leading to a net enhancement of the UV sensitivity from the ground to the sky. Our *in vivo* results support this perspective by corroborating the *ex vivo* findings in an intact eye. Intriguingly, *in vivo* and *ex vivo* measured RFs differ subtly. Whereas *ex vivo* RFs show a clear asymmetry peak at the horizon, *in vivo* RFs are more asymmetric across large proportions of the visual field (Fig. 5j). This is consistent with PC predictions since natural image patches located above the horizon tend to be vertically asymmetric (Fig. 1d) due to a gradient of stimulus power (Fig. 1b). Such RF pattern indicates that other mechanisms, apart from the S-opsin gradient, have to be involved. One possibility could be the circuitry mediating the asymmetric surround of J-RGCs ^33,40^, which is ventrally displaced and M-cone and rod-sensitive. Interestingly, the vertical gradient of stimulus power will flatten at lower ambient light levels, e.g., at dusk and dawn. In conjunction, the relative strength of UV sensitivity and the antagonistic surround would also decrease ^36,37^. In these conditions, efficient-coding hypothesis would predict a more homogeneous RF distribution across the dorsoventral axis. Conversely, the horizon will become more prominent at photopic conditions, where rods are less active. In such situations, the *in vivo* RF architecture should have a localized asymmetric streak, as measured in our *ex vivo* data. It would be revealing to test if the retina-wide RF organization is dynamically reshaped under scotopic and photopic conditions. Finally, to fully benefit from this panoramic retinal code, the eye should maintain a relatively constant position on the horizon. In agreement with this idea, eye and head movements stabilize the retinal image remarkably well, on average ~10 ° in azimuthal angle, during behavior ^35,43^.

A distinct, yet related question can be asked about the emergence of direction-selective (DS) computations in retinorecipient areas, such as the superior colliculus, where neurons integrate input from the entire retina, including the asymmetric streak. DS encoding can emerge as a consequence of an asymmetric and time-shifted surround, as shown before ^33,44^. Consistent with the measured center-surround asymmetry, some studies have described these neurons as sensitive to upward motion ^45^, whereas others do not find such specificity ^46,47^. The efficient coding interpretation, such as the one adopted here, suggests that the key to resolving this puzzle might be understanding the statistics of what the animal ought to see in nature.

Our theoretical predictions established qualitative links between properties of RFs and their elevation within the visual field. They can be therefore thought of as a first-order approximation of how the retina is adapted to large-scale, spatial statistics of natural scenes. The exact pattern of global retinal adaptation should vary across species occupying diverse environments. It has been found that dorsoventral opsin gradients are present in many different mammalian species. For example, the rabbit, Chilean subterranean rodent cururo, European mole, the shrew, and even the spotted hyena ^42,48^ show higher S-opsin expression in the ventral retina. However, not all ecological niches are identical. For example, in dense forests, the vertical gradients of luminance and contrast are less prominent, and a clear horizon line might not be apparent. Interestingly, forest mice species whose opsin distribution has been described present a spatially uniform opsin distribution ^49^. This further strengthens our hypothesis, which relates the global organization of the retina to the statistics of the ecological visual field. Understanding this adaptation in more detail will require a careful analysis of stimuli from the ecological sensory niche, as well as estimation of biophysical parameters such as biological SNR, RF size and tiling to refine our theoretical predictions. The combination of these approaches will be a critical requirement for building a more general theory of vision across the animal kingdom ^50^.

## Acknowledgments

We thank Hiroki Asari for sharing the dataset of naturalistic images, Anton Sumser for sharing visual stimulus code, Yoav Ben Simon for initial explorative work with the generation of AAVs, and Tomas Vega-Zuñiga for help with immunostainings. We also thank Gasper Tkacik and members of the Neuroethology group for their comments on the manuscript. This research was supported by the Scientific Service Units of IST Austria through resources provided by Scientific Computing, the Preclinical Facility, the Lab Support Facility, and the Imaging and Optics Facility. This work was supported by European Union Horizon 2020 Marie Skłodowska-Curie grant 665385 (DG), Austrian Science Fund (FWF) stand-alone grant P 34015 (WM), Human Frontiers Science Program LT000256/2018-L (AS), EMBO ALTF 1098-2017 (AS) and the European Research Council Starting Grant 756502 (MJ).

## Author contributions

DG, WM and MJ designed the study. DG and JS performed the retina imaging experiments with help from MJ. DG and MJ developed the retinal imaging system. JS and MJ developed the shifting stimuli. AS performed the *in vivo* imaging experiments and developed the imaging system with help from MJ. DG, AS and OS analyzed the experimental data with inputs from WM and MJ. WM analyzed natural scene statistics and developed the normative model. DG, WM and MJ wrote the manuscript with help of the other authors.

## Competing interests

Authors declare that they have no competing interests.

## Data availability

All data used in the analysis will be made available at IST DataRep upon publication.

## Code availability

All code used to generate the results will be made available at Github upon publication.

## Materials and Methods

### Theory

We modeled neural responses *r_t_* as a dot product of the model receptive field 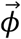 and noisy stimulus vectors (image patches) 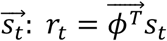, where *T* denotes vector transposition. Receptive fields (filters) were optimized to minimize the following cost function:

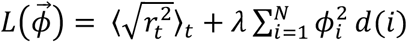

Where *d(i)* is the squared distance between the i-th value of the RF and the one with the peak absolute value, and *λ* is the strength of the spatial locality constraint. This form of the locality constraint has been introduced in ^11^, and it has been demonstrated that it is consistent with RGC properties ^11,13^. We note that the activity-related term in the cost function is equivalent to maximizing the sparsity of the neural activity quantified as the average absolute value of neural responses ^8^. Without the spatial locality constraint (i.e., *λ* = 0) the optimal RF is an oriented, Gabor-like filter. During optimization, to avoid convergence to trivial solutions, the norm of the RF 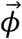 was constant and equal to 1. Overall, this cost-function enforces minimization of activity conveyed downstream, while preserving the information about the image and meeting the locality constraints. Conceptually, this goal is equivalent to that of the PC model ^10^. Our model generates predictions consistent with the PC model proposed in ^10^. (see Supplementary Fig. 1). It is, however, more flexible, and enables us to capture changes in the center size.

We modeled output of photoreceptor cells (stimuli *s_t_*) as natural image patches distorted with the additive Gaussian noise with variance *σ^2^* i.e.: *s_t,i_* = *x_t,i_* + *ξ*, where *ξ*~*N*(0, *σ*^2^) is the noise term. To simulate different SNR conditions, we manipulated the noise variance level, and optimized RFs for each of the noise levels separately.

We optimized RFs by numerically minimizing the cost function L via gradient descent. For training, we used a dataset of 50 000 square image patches 27 × 27 pixels in size taken from a dataset of natural images from the mouse visual environment ^18^. We sampled images uniformly across the upper and lower visual fields. Each image patch was normalized to have a zero mean and unit variance. To simulate the impact of changing SNR, we normalized images with added noise. During optimization, the dimensionality of natural image data was reduced with Principal Component Analysis (PCA) to 128 dimensions. For each noise level, dimensionality reduction was performed using the same matrix of PCA components computed on noiseless data. To simulate the impact of changing SNR homogeneity, prior to normalization we multiplied the bottom half of each image by a scaling factor lesser than one. After such scaling, we normalized the data and added noise of constant variance. During optimization of RFs on asymmetric stimuli, we computed PCA for each level of SNR asymmetry separately.

In all cases, prior to optimization, in order to enforce that the RF is centered in the image patch, we initialized RFs with random Gaussian noise with variance equal to 0.1 but set the central pixel value to −1. We independently optimized receptive fields using images taken in the ultraviolet and green parts of the spectrum, as well as in the “joint” spectrum where intensity of each pixel was average of green and ultraviolet values.

To evaluate RF properties, we defined the size of the RF to be the smallest circle which included 90% of energy (i.e. 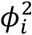) of optimal RFs averaged across all noise levels (Fig. 1 b, c).

Within that circle, we defined the center to be all 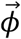 values smaller than 0, and the surround to be those larger than or equal to 0. The strength of the surround was thus equal to ∑_*i:ϕ_i_*≥0_|*ϕ_i_*| and the center to ∑_*i:ϕ_i_*<0_|*ϕ_i_*|. Sizes of center were simply numbers of entries that were smaller than 0.

To characterize changes in contrast and luminance across the visual field we used natural images published in ^18^. We limited our analysis to UV images only, however light statistics of the green channel do not differ qualitatively. The images provided in ^18^ were divided into two classes - upper visual field and lower visual field. To simulate the visual horizon, we concatenated pairs of images randomly selected from the upper and lower visual fields. We created a dataset of 1000 such concatenated images, and used them to compute the mean and variance of light intensity as estimates of local luminance and contrast, as well as to estimate the SNR as a function of elevation. To estimate the vertical asymmetry of the SNR pattern we used a square stimulus window, and a fixed noise variance. We note that key, qualitative aspects of our predictions do not depend on these choices. For each position *y* of the window along the vertical dimension of the visual field, we computed the vertical SNR asymmetry as: 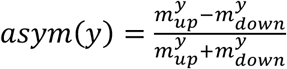 where 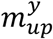 and 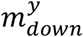 are sums of the SNR value within the stimulus window above and below its midline respectively.

### Animals

Mouse protocols were reviewed by the institutional preclinical core facility (PCF) at IST Austria. All breeding and experimentation were performed under a license approved by the Austrian Federal Ministry of Science and Research in accordance with the Austrian and EU animal laws. For retinal experiments, triple transgenic male and female mice, age 5-12 weeks were used for this study (Vglut2-ires-cre (JAX 28863), TITL-R-CaMP1.07-D (JAX 030217) and ROSA26-ZtTA (JAX 012266)). For *in vivo* imaging experiments, C57BL/6J (JAX 000664), aged 6-11 weeks at eye injection, were used. The mice were housed in a standard (*in vivo*: inverted) 12-hour day-night cycle and euthanized by cervical dislocation before *in vitro* imaging.

### *Ex vivo* imaging

The dark-adapted mouse retina was isolated under far-red light (LED peak 735 nm, additionally filtered with a 735 nm LP filter eliciting an isomerization rate of ~17 R*/s) in oxygenated Ames’ medium (Sigma) with constant bubbling (95% O2, 5% CO2) at room temperature. Left and right retinas were kept separate for identification. Four incisions were made to flat-mount the retina, with ganglion cells facing up, on a 18 mm coverslip (VWR 631-0153), and held down with a filter paper (Merck GSWP01300) having a ~2.5 mm x 2.5 mm imaging window cut out. The preparation was then placed in a heated (32 °C) superfusion chamber on the stage of a custom-built upright fluorescence microscope. The retina was left to recover for a minimum of 10 minutes with the excitation light of the microscope turned on. An amber LED (Thorlabs M595L4) filtered with a BP filter (Thorlabs FB580-10) was used for excitation and a BP filter (Thorlabs 641-75) in series with a 600nm LP filter (Thorlabs FEL0600) was used for collection. Background excitation light intensity was at a constant mean photopic intensity of 10^5^ R*s^−1^ per rod (at 585 +/− 5 nm). Isomerization rates were determined using opsin templates ^51^ and assuming that the mouse rod has an optical density at peak absorption wavelength of 0.015 μm^−1^, a length of 24 μm, a diameter of 1.4 μm and a quantum efficiency of 0.67 ^52,53^. Each retina was tiled by recording 3-7 different fields of view (FOV) at 10x magnification (Olympus XLUMPLFLN20XW objective) using a sCMOS camera (Photometrics Prime 95B) at 10 fps and 1.1 μm/pixel resolution.

### Visual stimuli for retinal experiments

Light stimuli were delivered from a modified Texas Instruments DLPLCR4500EVM DLP projector through a custom-made lens system and focused onto the photoreceptors (frame rate 60 Hz, magnification 2.5 μm per pixel, total area 3.2 mm x 2 mm). The projector’s blue LED was replaced with a high-power UV-LED (ProLight 1 W UV LED, peak 405 nm), to improve the differential stimulation of S pigments. Two SP filters in series (Thorlabs FESH0550) were put in the stimulus path to block green light from entering the camera. Intensities and spectra were measured using a calibrated spectrometer (Thorlabs CCS-100) and a digital power meter (Thorlabs S130C sensor). A shifting spatiotemporal white-noise stimulus was presented using a binary pseudo-random sequence, in which the two primary lights (green and UV) varied dependently. All white-noise stimuli were presented at 6 Hz update for 15 min. The checker size was 100 × 100 μm and the entire grid was shifted by random multiples of 10 μm in both x- and y-axis after every frame. In comparison experiments, static checkers (without shifts) of 100 × 100 μm and 25 × 25 μm were interleaved with the shifting checkers in chunks of 5 min for a total of 20 minutes for each of the three checker types. A ‘chirp’ stimulus with a 1s bright step, increasing amplitude (0 to 127 over 8 s) and increasing frequency (0 to 4 Hz over 8 s) was repeated for 5 trials to reproduce clustering of responses ^20^. Moving square gratings (0.6 cycles/s temporal frequency and 0.025 cycles/μm spatial frequency) or a wide bright bar (1 mm/s speed, 2 mm width) in 8 directions, repeated for 5 trials, were used for assessing direction selectivity. All visual stimuli were generated using the Psychtoolbox ^51^.

### Histology

After the *ex vivo* recordings, some of the retinas were fixed with 4 % PFA for 30 min and stained for S-opsin and RFP. Retinas were incubated for 7 days at 4 °C in PBS, containing 5 % donkey serum, 0.5 % Triton X-100, goat anti S-opsin (1:500, Rockland 600-101-MP7) and rabbit anti-RFP (1:1000, Rockland 600-401-379). After washing thrice in PBS for 15 min each, retinas were incubated overnight in secondary antibodies, donkey anti-goat Alexa Fluor 488 (1:1000 Abcam ab150129) and donkey anti-rabbit Alexa Fluor 594 (1:1000, Invitrogen R37119). Retinas were then mounted and imaged with Olympus VS120 Slidescanner with a 20x objective. For cell-type characterization, Vglut2-ires-cre; TITL-R-CaMP1.07-D; ROSA26-ZtTA mice were euthanized and perfused intra-cardially, followed by retina extraction and staining for RBMPS or SMI-32, along with RFP (primary antibodies: guinea pig anti-RBPMS (1:500, Sigma ABN1376), mouse anti-SMI32 (1:500, BioLegend 801701), rabbit anti-RFP (1:1000, Rockland 600-401-379) or mouse anti-RFP (1:500, MBL M155-3); secondary antibodies: goat anti-Guinea pig Alexa Fluor 647 (Invitrogen A21450), donkey antimouse Alexa Fluor 647 (1:1000, Abcam A-31571) and donkey anti-rabbit Alexa Fluor 594 (1:1000, Invitrogen R37119). The staining protocol was the same as above and these retinas were imaged with Leica SP8 confocal microscope.

After the final *in vivo* recording, mice were terminally anesthetized with Ketamine/Xylazine (100 mg kg^−1^, 10 mg kg^−1^) i.p. and transcardially perfused with PBS, followed by 4 % PFA. Brains were extracted and post-fixed in 4 % PFA at 4 °C overnight. Brains were then washed and transferred to 30% sucrose solution for cryoprotection overnight at 4°C and subsequently frozen and the midbrain coronally sliced into 40 um sections on a Leica SM2010R sliding microtome. Sections were washed and then incubated in PBS, containing 5 % donkey serum, 0.3 % Triton X-100, goat anti GFP (1:2000, Abcam, ab6673) overnight at 4°C. After washing thrice in PBS for 15 min each, brain sections were incubated for 1 hour in secondary antibody solution, donkey anti-goat Alexa Fluor 488 (1:1000 Abcam, ab150129), washed thrice again in PBS and mounted on slides, where they were stained with DAPI (not shown) and coverslipped with custom-made mowiol. Brain sections were imaged with Nikon CSU-W1 spinning disk confocal microscope at 20x tile stack acquisition. Image stacks were shading corrected with BaSiC ^52^ in ImageJ and finally maximum-projected over the whole stack.

### Preprocessing

ROIs were detected automatically from raw calcium movies using Suite2p ^54^. Fluorescence traces, *F*, were detrended by computing *ΔF/F*, where the 8^th^ percentile in a 20 s sliding window centered around each time point was taken as the baseline fluorescence. Different fields of view from the same retina were stitched together based on coordinates from the stage motors and repeated ROIs in overlapping regions were manually annotated using an open-source tool ^53^. Repeated ROIs with the highest score in Suite2p’s built-in classifier were kept for analysis. The deconvolved signal from Suite2p (with *tau* = 1.0 s) was used for calculating receptive fields.

The median *ΔF/F* response across trials was taken as the response to the chirp and normalized by dividing by the maximum of the absolute values across time. Quality Index was computed as in ^20^, and only responses with QI > 0.45 were kept for clustering (8019 out of 30798 neurons). For moving gratings and bar, the mean across trials and maximum across time was taken as the response in any one direction.

### Receptive Field Mapping

The receptive field for each neuron was computed as a calcium-triggered-average. The spatiotemporal receptive field at latency *τ,* position *(x, y)* for neuron *i* was computed as,

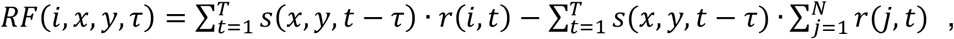

where *r(i,t)* is the deconvolved response of neuron *i* at time *t*, *s*(*x,y,t*) is the white noise stimulus, *T* is the length of the recording and *N* is the total number of neurons in the recording. The second term in this equation subtracts away the residual distribution of the stimulus and the contribution of slow bleaching that is common to all neurons in that recording and leads to receptive fields that had noticeably less noise. Only the UV channel of the stimulus was used for RF mapping. The latency *τ* was varied in increments of 0.025 s (40 Hz), and the stimulus was interpolated by using the last frame before a particular time, *t* – *τ*. The RF for each neuron was normalized between [-1,1] by subtracting the mean value of the RF at latencies *τ* < 0, and dividing by the maximum absolute value of the entire RF.

The location of the center of an RF was estimated by finding the pixel that varied the most across time, *P_var_* = *argmax Var^t^*(*x,y*) where *Var^t^*(*x,y*) is the variance across time for position (x,y). Each neuron’s RF was cropped within a square window of edge 1mm centered on this pixel. *SNR* of an RF was computed as the peak-to-noise ratio where the power of noise was estimated in regions with distance >0.5mm from the point *P_var_*. Only RFs with peak SNR > 15dB were kept for analysis (31135 selected RFs out of 37086 recorded neurons). The location of the RF in time was found in a similar way; *T_var_* = *argmax Var^xy^* (*t*) here *Var^xy^*(*t*) is the variance in space.

### Parametrization of Receptive Fields

We parametrize spatiotemporal properties of the center and surround of the RFs as a sum of two 2-d Gaussian distributions (Gaussians) 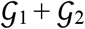. The first Gaussian represents the center of the RF; its amplitude can be either negative or positive corresponding to an OFF or an ON cell. The amplitude of the second Gaussian is required to be of the opposite sign to model the properties of the surround. We compute a spatial representation of the RF, denoted as RF2D as the median of the RF within a small time window around *T_var_*. To reduce noise, we exclude the pixels that are weakly correlated with *P_var_* across time. The sum of 2-d Gaussians 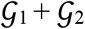 is then fitted to RF_2D_. Each Gaussian is defined by the amplitude *A*, center (*m^x^,m^y^*), width (*σ^x^,σ^y^*) and the orientation angle *θ*. We start by fitting only one Gaussian 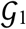 to parametrize the location (*m^x^, m^y^*), the size (*σ^x0^, σ^y0^)*, and the orientation of the center. In the next step we fit the sum of Gaussians, where we fix (*m^x^*, *m^y^*, *σ*^*x*0^, *σ*^*y*0^, *θ*_1_) parameters of 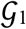 while allowing all other parameters to be fitted anew.

Here we differentiate two types of RFs: a RF with a strong center and a weaker surround that largely overlap, and a RF where both center and surround components are strong and well separated. For the first case, we impose a constraint such that the center of 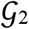 is within the distance 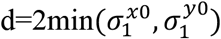 from the center 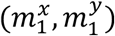. We implement this constraint as a penalty sigmoid function 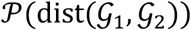 of the distance between the locations of the center and surround components. We add 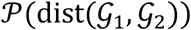 to the Gaussian mixture and allow it to be prohibitively large for dist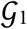. Encoding the constraint in this manner allows us to remove the bias for the location of the surround on the diagonals of the RF_2D_ which otherwise happens when fitting Gaussian mixture on the square bound-constrained region. There are no constraints for the second type of the RFs where the surround component is strong and more distant from the center. To decide the type of the RF, we find the maximum and minimum points of the RF, and we compute the distance between them and the ratio of their absolute values. If the ratio of the smaller to the bigger values is less than 0.75 or if the distance between the extrema points is less than *d,* then we classify such a RF as the first type, and as the second type otherwise. Experimentally we found that imposing such a constraint on the location of the second Gaussian leads to a better fitted sum of Gaussians for RFs with largely overlapping center and surround components. All the fitting procedures were implemented using the nonlinear least-squares solver `Isqcurvefit` in MATLAB.

Using the parametrization, we can compute various RF characteristics. We find two sets of pixels corresponding to the center and the surround. Center pixels are the pixels within two standard deviations from the center of the 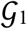. Surround pixels are the pixels within two standard deviations from the center of the 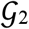 and that are not center pixels. The *center size* is the number of pixels in the center set, converted to mm^2^ for display. The *relative surround strength* is the ratio of the absolute value of sum of surround pixels to the absolute value of sum of center pixels. *Vertical surround asymmetry* is defined as (*u* – *l*)/(*u* + *l*) where *u* and *l* denote the absolute value of sum of pixels in the upper and lower halves of the surround pixels, respectively. The *distance* between the center and the surround is the distance between the center of mass (COM) of the center pixels and the COM of the surround pixels. The *orientation* is the angle between the horizontal axis and the line connecting the two COMs. *Radial profiles* were computed as the average values of the pixels in RF_2D_ within rings of increasing radii centered on the point *P_var_*. The average value of the center or surround pixels across time was taken to be the *RF center or surround temporal dynamics,* respectively. The *R2 goodness of fit* is computed as 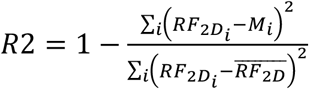, where 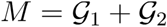 is the RF parametrization. The values of the above RF parameters are reported for a few representative neurons in Supplementary Fig. 5.

### Retina alignment

All functional imaging experiments were performed with randomized retina orientations. Retinas that were co-stained against S-opsin and RCaMP1.07 (n = 6 retinas) and the direction with the highest density of S-opsin was taken to be the ventral direction ^54^. The stitched maximal projection images of the functional imaging experiments were aligned to the RCaMP channel using the retinal vasculature. In each of these retinas, we observed that a streak of asymmetric surrounds was always consistently present across the dorsal retina. Thus, we assumed this to be a reproducible feature, which we then used to manually align the remaining n = 5 retinas that did not have an S-opsin staining (and hence no ground truth orientation). To avoid any potential circular arguments, we present the location of neurons from these 5 retinas only in Supplementary Fig. 8, with a clear indication that the retinal orientation is presumed.

The coordinates of cells from each retina were then translated to make the optic nerve the zero of the coordinate system and rotated such that the positive y-axis denoted ventral direction. Left retinas were flipped in the nasotemporal axis such that the positive x-axis denoted nasal direction for all retinas. All spatial receptive fields were also translated and rotated accordingly. The cartesian retinal coordinates of cells in the stained retinas were converted to spherical visual coordinates using the R-package Retistruct, assuming the optical axis of the mouse eye to be oriented at 64 deg azimuth and 22 deg elevation from the long axis of the animal ^31^.

### Binning of RF properties

For 2-d bins, the retinal space from −1500 μm to 1500 μm in both nasotemporal (NT) and dorsoventral (DV) axes was divided into a square grid and all neurons within each bin were collected. Only 2-d bins with at least 5 neurons were analyzed to minimize sampling bias. The spatial RF of all neurons within each bin of size 300 μm were averaged and plotted at the location of the bin in (Fig. 3a, Supplementary Fig. 7). The RF parameter values of neurons within each bin of size 50 μm were averaged to yield a 2-d map of the parameter, and this map was visualized (without smoothing) in (Fig. 3d-h). Owing to its area-preserving property, the sinusoidal projection of visual space was binned in the same way as the retinal surface, and the fraction of cells in each bin that had ventrally oriented surrounds were plotted in Fig. 3k.

For 1-d analysis, the bins were defined along the DV or NT axis based on the range of coordinates in a particular group (by retina (Fig. 3) or cluster (Fig. 4)). The range was divided into 6 equally spaced bins and the mean parameter value of neurons within each bin was plotted at the coordinate of the center of the bin. As a summary statistic, two-sample Kolmogorov-Smirnov-test was performed between ventral and dorsal samples of these binned values (n = 56 samples for Fig. 3 and n = 60 samples for Fig. 4). Two-sample Kolmogorov-Smirnov tests (with Bonferroni correction for multiple comparisons) were also performed between raw values of RF parameters for Supplementary Fig. 7. In addition, the weights of linear regressions of RF parameters in elevation (dorsoventral) and azimuth (nasotemporal) orientations are reported.

### Clustering into functional types

The Gaussian Mixture Model procedure developed in ^20^ was used for clustering temporal RFs and chirp responses, separately. In brief, after normalization, the trace was first reduced in dimension using Principal Component Analysis (10 components for temporal RFs and 20 components for chirp responses) and then GMM models with diagonal covariance matrices were fitted while increasing the number of clusters. The numbers of clusters were identified to be 10 for temporal RFs and 20 for chirp responses based on elbow points in the respective Bayesian Information Criteria (BIC) curves. One chirp cluster (n = 68 cells) lacked stimulus-evoked responses and was discarded on visual inspection. To assess the degree of overlap between the RFs of neurons belonging to each of the clusters, we defined the tiling index of the cluster *K* as the area of the union of all RF centers in a cluster, divided by the sum of their individual areas:

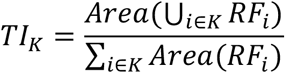

The value of this index for each cluster was computed separately for each retina and the mean and STD across retinas are reported in Supplementary Table 1.

### Viral eye injections

For viral-mediated gene transfer, 6-11week old wildtype C57BL/6J mice (JAX 000664) were anesthetized with ketamine/xylazine by intraperitoneal injection. A 30 1/2G needle was used to make a small hole in the temporal eye, below the cornea. 1 μl of vitreous fluid was withdrawn and then 1 μl of AAV2.7M8-syn-GCaMP8m viral vector solution (at a titer of ~1 × 10^13^ genome copies per ml, ISTA viral facility) was injected into the subretinal space with a Hamilton syringe and 33G blunt-ended needle.

### Mouse surgery for *in vivo* imaging

2-3 weeks after viral eye injections mice were injected with meloxicam (20 mg kg^−1^ s.c., 3.125 mg ml^−1^ solution) and dexamethasone (0.2 mg kg^−1^ i.p., 0.02 mg ml^−1^ solution). Anesthesia was induced by 2.5 % Isoflurane in oxygen in an anesthesia chamber. The mouse was subsequently fixed in a stereotaxic device (Kopf) with a constant isoflurane supply at 0.7 to 1.2 % in O_2_ and body temperature controlled by a heating pad to 37.5 °C. After the assertion that reflexes subsided, the cranium was exposed and cleaned of periost and connective tissue. A circular craniotomy of 4 mm diameter was drilled above the left superior colliculus and from this point onwards, the exposed brain was constantly irrigated with artificial cerebrospinal fluid. The exposed dura mater was removed, subsequently, the left transverse sinus was sutured twice with 9-0 monofil surgical suture material (B. Braun) and cut between the sutures. Cortical areas covering the left superior colliculus were aspirated with a cell culture vacuum pump (Accuris) connected to a blunt needle with 0.5 mm diameter. A 3 mm circular coverslip was glued (Norland optical adhesives 61) to a thin-walled custom-made conical ring, made from stainless steel. The coverslip-ring was inserted into the cavity left by the aspirated cortex, so that the glass sits flush on the surface of the superior colliculus. Slight pressure was applied with the help of a thinned toothpick, fixed to the stereotaxic arm. The space around the insert was filled with Dura-Gel (Cambridge Neurotech) and the insert was fixed in place with VetBond (3M). Then after cleaning and drying the surrounding cranium, a multilayer of glues was applied. First, to provide adhesion to the bone, All-in-One Optibond (Kerr) was applied and hardened by blue light (B.A. Optima 10). Second, Charisma Flow (Kulzer) was applied to cover the exposed bone and fix the metal ring in place by also applying blue light. After removal of the fixation toothpick, a custom-designed and manufactured (RPD, Vienna) headplate, selective laser-sintered from the medical alloy TiAl6V4 (containing a small bath chamber and microridges for repeatable fixation in the setup), was positioned in place and glued to the Charisma on the cranium with Paladur (Kulzer). Mice were given 300 μl of saline and 20 mg kg^−1^ meloxicam (s.c.), before removing them from the stereotaxic frame and letting them wake up while kept warm on a heating pad. Another dose of 20 mg kg-1 meloxicam s.c. and 0.2 mg kg^−1^ i.p. dexamethasone was further injected 24 hours after conclusion of the surgery. After the implantation surgery animals were let to recover for 1 week.

### *In vivo* visual stimulation and eye movements

Mice were head-fixed while awake using a custom-manufactured clamp, connected to a 3-axis motorized stage (8MT167-25LS, Standa). Mice could run freely on a custom-designed spherical treadmill (20 cm diameter). Visual stimuli were projected by a modified LightCrafter (Texas Instruments) at 60 Hz, reflected by a quarter-sphere mirror (Modulor) below the mouse and presented on a custom-made spherical dome (80 cm diameter) with the mouse’s head at its center. The green and blue LEDs in the projector were replaced by cyan (LZ1-00DB00-0100, Osram) and UV (LZ1-00UB00-01U6, Osram) LEDs respectively. A double band-pass filter (387/480 HD Dualband Filter, Semrock) was positioned in front of the projector to not contaminate the imaging. The reflected red channel of the projector was captured by a transimpedance photo-amplifier (PDA36A2, Thorlabs) and digitized for synchronization. Cyan and UV LED powers were adjusted so that the reflectance on the screen matches the relative excitation of M- and S-cones during an overcast day, determined and calibrated using opsin templates ^55^ and a spectrometer (CCS-100, Thorlabs). Stimuli were designed and presented with Psychtoolbox ^51^, running on MATLAB 2020b (MathWorks, Inc.). Stimulus frames were morphed on the GPU using a customized projection map and an OpenGL shader to counteract the distortions resulting from the spherical mirror and dome. The dome setup allows the presentation of mesopic stimuli from ca. 100 ° on the left to ca. 135 ° on the right in azimuth and from ca. 50 ° below to ca. 50 ° above the equator in elevation.

Visual stimuli were similar to *ex vivo* retinal imaging experiments: A shifting spatiotemporal white-noise stimulus was presented using a binary pseudo-random sequence, in which the two primary lights (Cyan and UV) varied dependently. All pseudo white-noise stimuli were presented at 5 Hz update in 5 min episodes, interleaved with different stimuli (e.g., grey screen, moving gratings (not shown)) with a total pseudo white-noise duration of 15-60 (median = 25) minutes per recording. The checker size was 8 × 8 ° visual angle and the entire grid was shifted by random multiples of 0.4 ° visual angle in both elevation and azimuth axis after every frame. Eye movements of the right eye were recorded with a camera (Basler acA1920-150um, 18-108 mm macro zoom lens (MVL7000, ThorLabs), set at 100 mm, and infrared illumination 830 nm) via an infrared mirror at 50 fps.

### *In vivo* retinal terminal imaging

Two-photon axonal terminal imaging was performed on a custom-built microscope, controlled by Scanimage (Vidrio Technologies) running on MATLAB 2020b (MathWorks, Inc.) and a PXI system (National Instruments). The beam from a pulsed Ti:Sapphire laser (Mai-Tai DeepSee, Spectra-Physics) was scanned by a galvanometric-resonant (8 kHz) mirror combination (Cambridge Scientific) and expanded to underfill the back-aperture of the objective (16× 0.8 N.A. water-immersion, Nikon); 1.9 by 1.9 mm field-of-view; 30 Hz frame rates. Fast volumetric imaging was acquired with a piezo actuator (P-725.4CA, Physik Instrumente). Emitted light was collected (FF775-Di01, Semrock), split (580 nm long-pass, FF580-FDi01, Semrock), band-pass filtered (green: FF03-525/50; red: FF01-641/75, Semrock), measured (GaAsP photomultiplier tubes, H10770B-40, Hamamatsu), amplified (TIA60, Thorlabs) and digitized (PXIe-7961R NI FlexRIO FPGA, NI 5734 16-bit, National Instruments). Laser wavelength was set between 920 and 950 nm. Average laser output power at the objective ranged from 57 to 101 (median: 69) mW ^56^. A field-of-view of 0.32 - 1.85 (median: 0.68) mm^2^ was imaged over 3 – 7 (median: 6) planes with 14 – 40 (median: 25) μm plane distance at a pixel size of 0.6 – 1.9 (median: 1.3) μm and a volume rate of 4.3 – 9.5 (median: 5.0) Hz. Each mouse was recorded in 2-4 imaging sessions on different days. In a subset of mice (n = 2) in separate imaging sessions, absence of substantial z-motion was verified by injecting 40 μl of Dextran TexasRed (3000MW, X mg/ml, ThemoFisher Scientific) s.c. and imaging brightly red labeled blood vessels at 980 nm ^56^.

### *In vivo* eye movement analysis

Behavior videos were analyzed with DeepLabCut ^57^, labeling 8 points around the pupil. The 8 points were then fitted to an ellipse and the ellipse center position transformed to rotational coordinates under the assumption of eyeball radius = 1.5 mm ^58^, using custom python scripts. The median of all eye positions was set to zero azimuth and elevation, i.e., all eye coordinates are relative to the median position. The individual horizontal axis, which varied slightly between mice due to differences in the positioning of the head plate, was corrected by leveraging a behavioral feature of head-fixed mice: Saccadic movements are nearly exclusively in one plane ^35^. Saccades were extracted by determining events of fast position changes on a median filtered position trace (median filter window: 0.7 s, minimal saccadic speed: 45 °/s, minimal saccade amplitude: 3 °, minimal saccade interval: 0.25 s). The preferred saccadic orientation and orientation tuning was determined in a similar fashion as for neuronal visual orientation tuning based on circular variance:

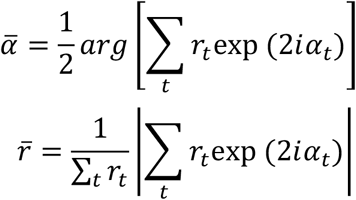

with 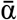 as the saccadic orientation angle, 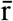 as saccadic orientation tuning, r_t_ as saccade amplitude and *α_t_* as direction of saccade t (Supplementary Fig. 10). Saccade orientation tuning was very high, with mean selectivity = 0.8.

### *In vivo* axonal terminal analysis

Functional calcium imaging data was first analyzed with suite2p (v0.10.0) ^59^ for motion correction and ROI extraction. ROIs were then curated manually based on morphological and activity shape. Further analysis was performed in custom MATLAB R2021a (MathWorks, Inc.) scripts: dF/F0 was estimated based on published procedures ^60^ by first subtracting neuropil contamination (from suite2p, fluorescence signal of 350 pixels surrounding the ROI, excluding other ROIs) with a factor of 0.5 (estimated from fluorescence of small capillaries as reported previously). From the neuropil-corrected ROI fluorescence, baseline F0 was defined as the 8th percentile of a moving window of 15 seconds ^61^. dF/F0 was then calculated by first subtracting and then dividing fluorescence trace by median of the same 15-second window ^60^. Fluorescence signal-to-noise ratio (fSNR) was defined for each neuron by dividing the 99th percentile of the dF/F trace (“signal”) by the standard deviation of its negative values after baseline correction (“noise”). Only axonal segments with fSNR ≥ 5 were included in further analysis. The deconvolved signal from Suite2p (with *tau* = 0.7 s) was used for calculating receptive fields. Note that multiple axonal ROIs can originate from the same retinal ganglion cell. Spatiotemporal receptive field analysis for *in vivo* retinal terminals was conducted as for *ex vivo* retinal ganglion cell imaging, but on visual stimuli downsampled to a resolution of 1 ° visual angle. The resulting 50 × 50 ° receptive fields are contaminated by eye movements and exhibit lower SNR (as determined by temporal variance of the most varying pixel over the temporal variance of pixels with > 50 ° visual angle distance) than *ex vivo* soma recordings, requiring further inclusion criteria: SNR > 15 (as in *ex vivo* data) and peak variance over time located at tau values between −0.1 and 0.6 s. Additionally, the retinotopic projection pattern of retinal ganglion cells to the superior colliculus was utilized by fitting a map from visual coordinates to collicular space. For each recording, receptive field center azimuth and elevation values were fitted separately to the location of the ROI in the superior colliculus using the “poly22” fit option in MATLAB and using only the highest 15 % ROI’s in SNR and SNR as a fitting weight. Boutons with peak location of the RF deviating more than 20 ° visual angle from its expected location based on the retinotopy fits were removed from further analysis (removing 828 boutons).

The main saccadic axis was used to rotate the computed spatial receptive fields around their respective center and the center positions in visual space as spherical rotation around the approximate eye axis (65 ° from frontal direction in horizontal plane). Finally, while freely moving mice hold their head at an approximate pitch angle of 30 ° downwards ^62^, *in vivo* imaging allowed only for a pitch angle of 10 ° downwards. To compensate, the center positions of all RFs were spherically rotated 20 ° downwards around the main pitch axis (90° from frontal direction in horizontal plane). Please note that these calculations only allow an estimate of the position of the horizon in free locomotion.

To avoid biasing the analyses by eye movements, receptive field parameterization was conducted on mean vertical 1-d profiles of extracted 50 × 50 ° 2-d RF crops at the peak azimuthal position ± 1 °, where ON center bouton RFs were inverted. To extract parameters from 1-d RFs they were fitted with a difference of 2 Gaussians, initialized with the central peak magnitude (Mpeak) and width (Wpeak). The fitting procedure was then constrained with amplitude_center_ ∈ [M_peak_/2, inf], amplitude_surround_ ∈ [0,inf], location_center_ ∈ [−20, 20]°, location_surround_ ∈ [−25, 25]° (edge of crop), sigma_center_ ∈ [W_peak_/4, inf] and sigma_surround_ ∈ [W_peak_, inf]. Boutons with center fit location in 1-d RFs more than 5 ° or with surround fit location more than 25 ° distant from peak estimation based on variance in 2-d RFs, were excluded from further analysis (1609 boutons removed). Extraction of parameters was identical to *ex vivo* RF parameterization, except center size, where *in vivo* 2sigma_center_ of the fit was used.

For presenting RF characteristics of RGC axonal boutons in the SC, the centered 1-d RFs were binned and averaged in each bin. RF parameters varying over elevation and azimuth are presented as parameters of the fit on the mean 1-d RF in the respective bin. Linear regression weights were computed from the parameters and location of each individual bouton.

## Supplementary Material

### Model Details

Here we follow the derivation and assumptions proposed in ^10^. The receptive field model in ^10^ assumes that the center pixel *s_t,cent_* of the t-th stimulus 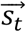 is subtracted from its linear prediction computed from the surround 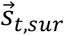. Instead of encoding the raw value of the central pixel, the model RGC encodes the difference between this prediction and the center in order to minimize the dynamic range of its output. The optimal prediction weights 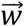 are optimized to minimize the mean squared error:

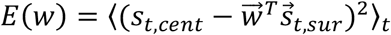

where *T* denotes vector transposition.

The optimal vector of surround weights 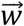 is a solution to the following equation:

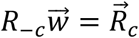

where *R_i,j_* = 〈*s_t,i_s_t,j_*〉_*t*_ is the spatial autocorrelation of natural images, and *i,j* index pixels within an image patch, 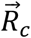 is the autocorrelation vector of the center pixel with all other pixels, and *R_-c_* is the square correlation matrix of all pixels without the center pixel.

The correlation function *R_i,j_* is approximated analytically as:

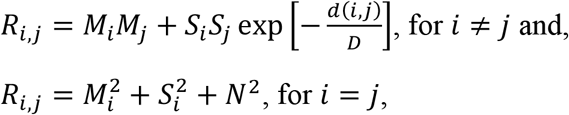

Where *M_i_* and *S_i_* are mean and standard deviation of the i-th entry of the image intensity respectively, *d(i,j)* is the Euclidean, spatial distance between entries labeled *i* and *j, D* is a constant controlling the decay of the correlation, and *N* is the standard deviation of the noise. The term *N_i_N_j_* vanishes for *i* ≠ *j* because noise is assumed to be uncorrelated.

To approximate the spatial autocorrelation function of natural images as a function of elevation within the visual field, we created a dataset of images with simulated horizon as described in the Methods. We then divided each of these images into uniformly separated horizontal bands. We sampled square image patches within each band. We then computed mean vectors 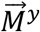 and standard deviation vectors 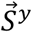, where the upper index *y* indicates the elevation band. Individual entries of these vectors corresponded to mean and variance of pixel values within the elevation band *y* respectively. We assumed constant values of the decay constant *D* and noise standard deviation *N*. Ratio of surround-to-center strength and surround asymmetry were computed as described in the Methods. We note that our results do not depend qualitatively on parameter choice and reveal similar trends across a broad range of parameter values.

## Supplementary Figures and Tables

**Supplementary Fig. 1.**
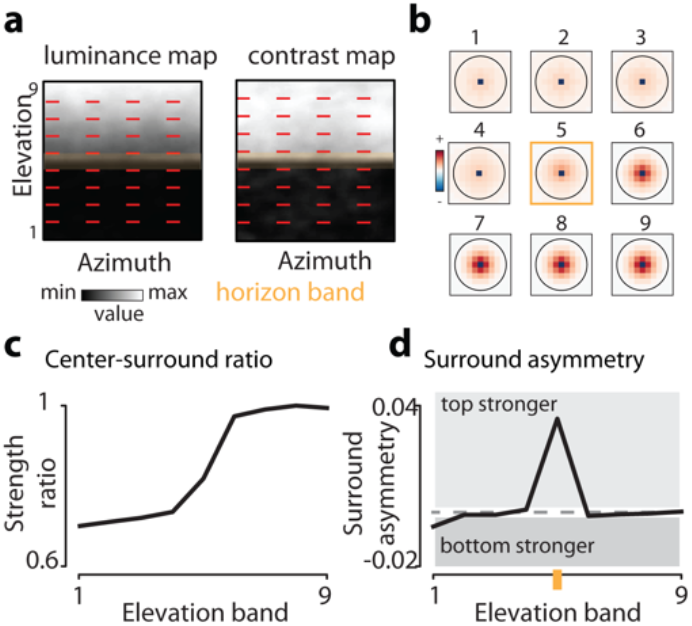
Predictive coding predictions derived from a previously proposed model. (**a**) Average luminance (left panel) and local contrasts (right panel) in the mouse field of view in the ultraviolet range. Red dashed lines separate elevation bands. Orange opaque rectangle denotes the horizon band. Natural image data - courtesy of Hiroki Asari ^18^. (**b**) Receptive fields optimized for each band in (a). (**c**) Relative surround-to-center strength as a function of the elevation band. (**d**) Surround asymmetry as a function of elevation band. Orange mark denotes the horizon band.

**Supplementary Fig. 2.**
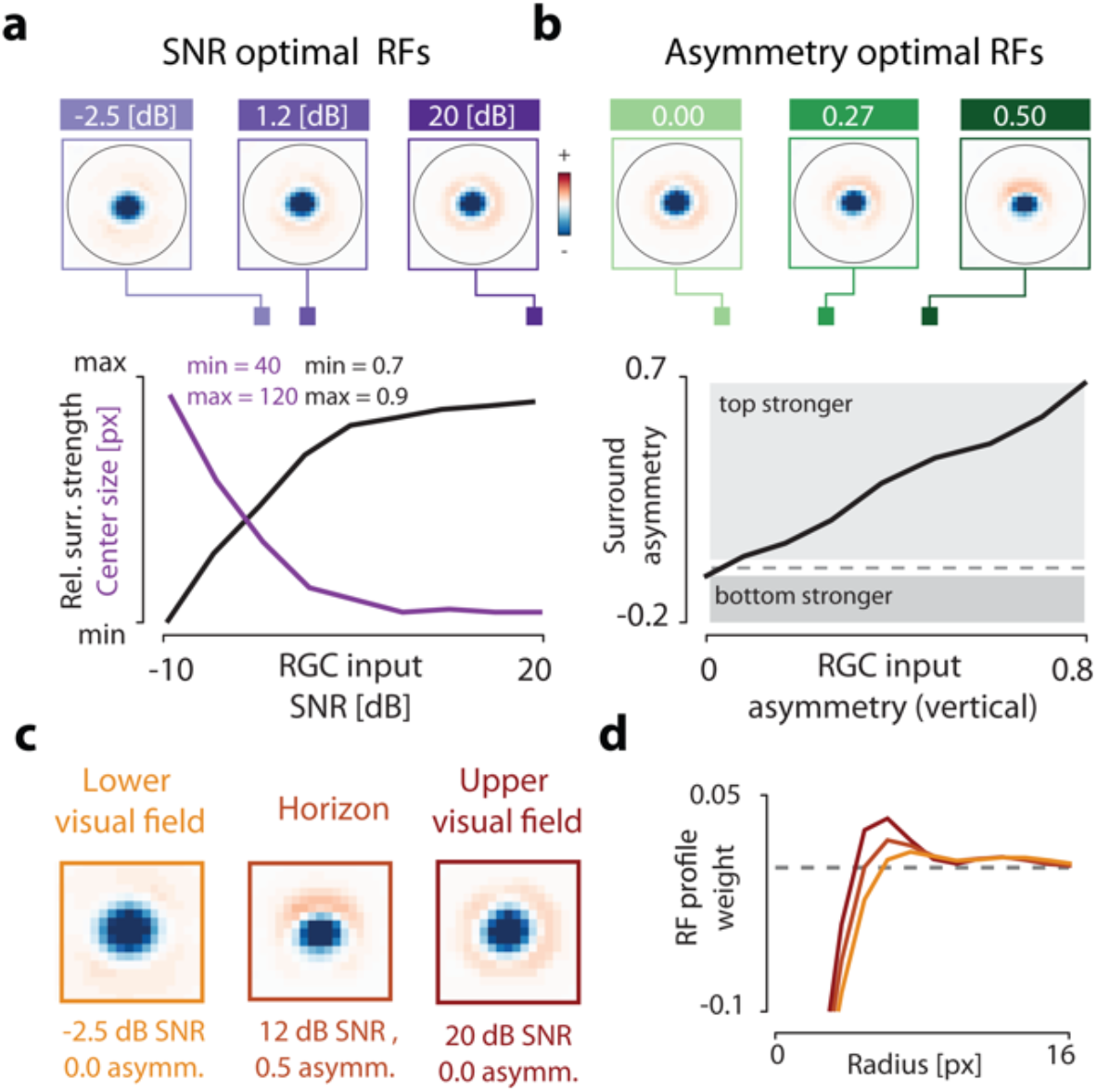
Predictive coding model trained with an alternative set of natural images. We used natural images of animals and landscapes of the African savannah. (**a**) Top - RFs optimized at different levels of SNR. Bottom – center size (purple line) and relative surround strength (black line) plotted as a function of the SNR. (**b**) Top - RFs optimized at different levels of the vertical SNR asymmetry. Bottom – vertical surround asymmetry plotted as a function of the vertical SNR asymmetry of the model photoreceptor output (RGC input). (**c**) RFs predicted for different positions within the visual field. (**d**) Horizontal cross-sections of model RFs in (c).

**Supplementary Fig. 3.**
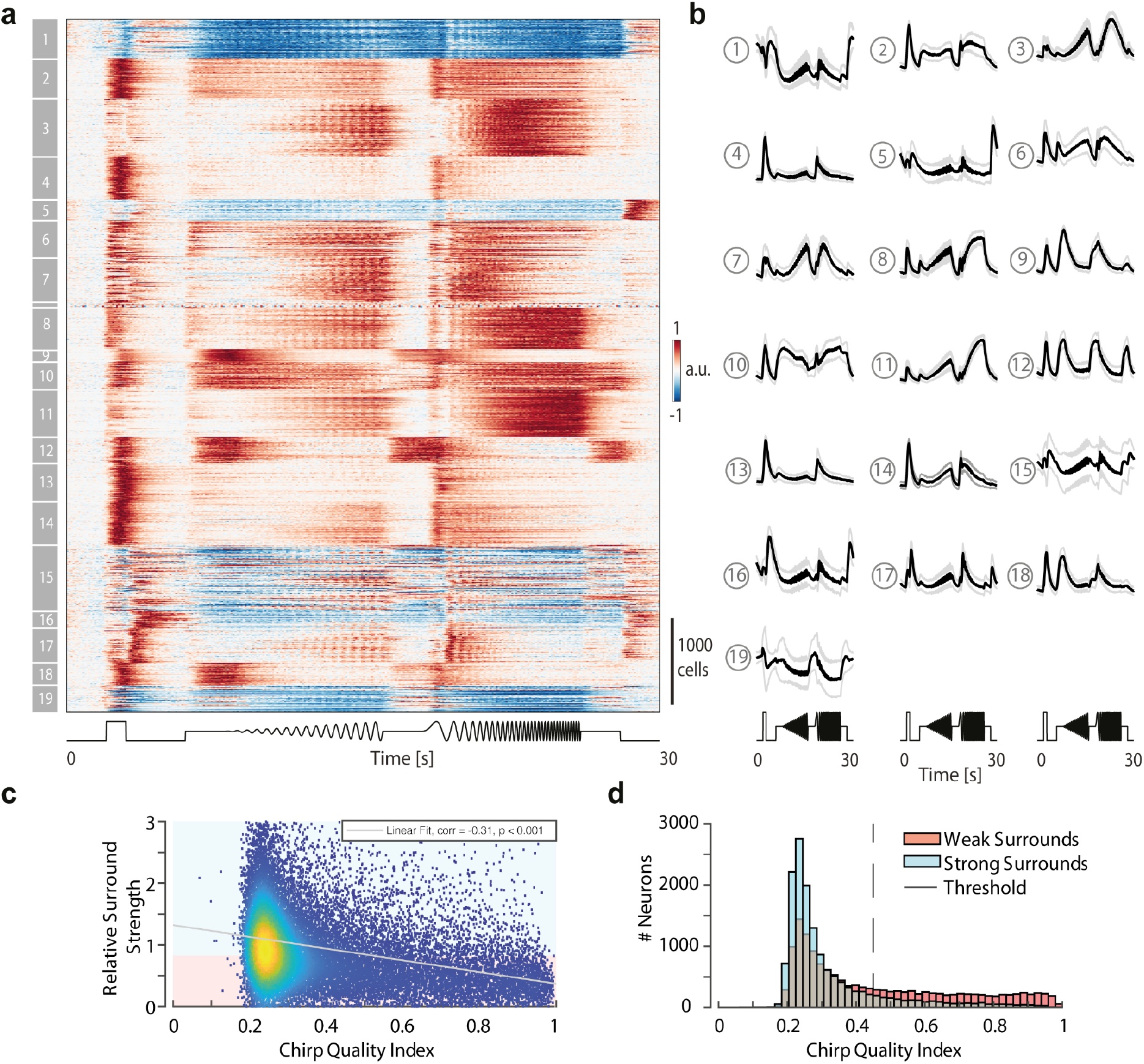
Clustering of Ca^2+^ signals to “chirp” stimulus shows segregation into functional types. (**a**) normalized Ca^2+^ responses to changes in frequency, contrast, and luminance, known as “chirp” stimulus from RGCs, sorted by cluster ids, as determined by GMM. (**b**) Mean and standard deviation of each cluster reveal ON (cluster 2, 4, 8, 9, 10, 11, 13, 14, 18), OFF (5, 12, 15, 16), ON-OFF (3, 6, 7, 17), suppressed-by contrast (1, 19) RGC-types, as well as differences in frequency tuning to slow (e.g., cluster 4), mid (e.g., cluster 3), fast (e.g., cluster 11) and all frequency modulations (e.g., cluster 2) or difference in luminance sensitivity (e.g., compare ON responsive cluster 14 and 18). Note: clusters 15 and 19 are not homogeneous, as seen in their large standard deviations. (**c**) Chirp-response quality index versus surround strength, determined using their spatiotemporal filters (as in Fig. 2J). Good-quality chirp responses have a strong bias for weak surrounds. (**d**) Distribution of chirp quality index for weak and strong surround, as defined in (c, see color code).

**Supplementary Fig. 4.**
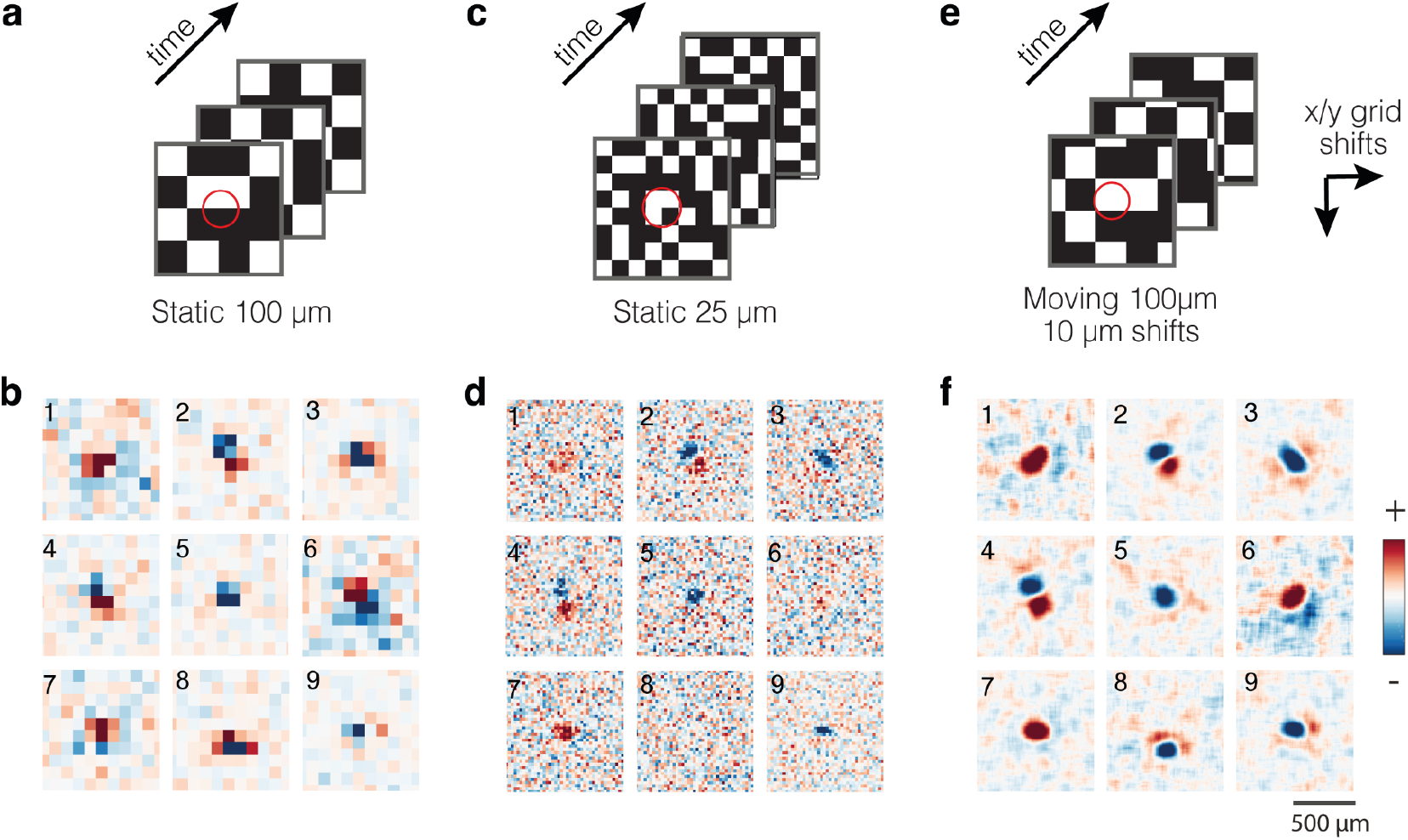
Comparison of RFs recovered from three different white-noise stimuli. (**a**) Schematic of binary white noise stimuli used for recovering RFs during imaging experiments for static checkers with grid size 100 × 100 μm^2^. (**b**) Spatial RFs of 9 representative neurons were generated from the stimuli above. (**c,d**) Same as (a,b) for the same neurons, but for static 25 μm sized checkers. (**e,f**) Same as in (a,b) for the same neurons but for a shifting checkers with grid size 100 × 100 μm^2^, where the entire grid was shifted by random multiples of 10 μm in both x- and y-axis. While the static 100 μm checkers were too low resolution for automatic analysis and the static 25 μm checkers were unable to drive many neurons strongly enough to elicit sufficient responses for RF reconstruction, the moving white noise stimulus was able to unambiguously recover the most detail in the center-surround structure of RFs.

**Supplementary Fig. 5.**
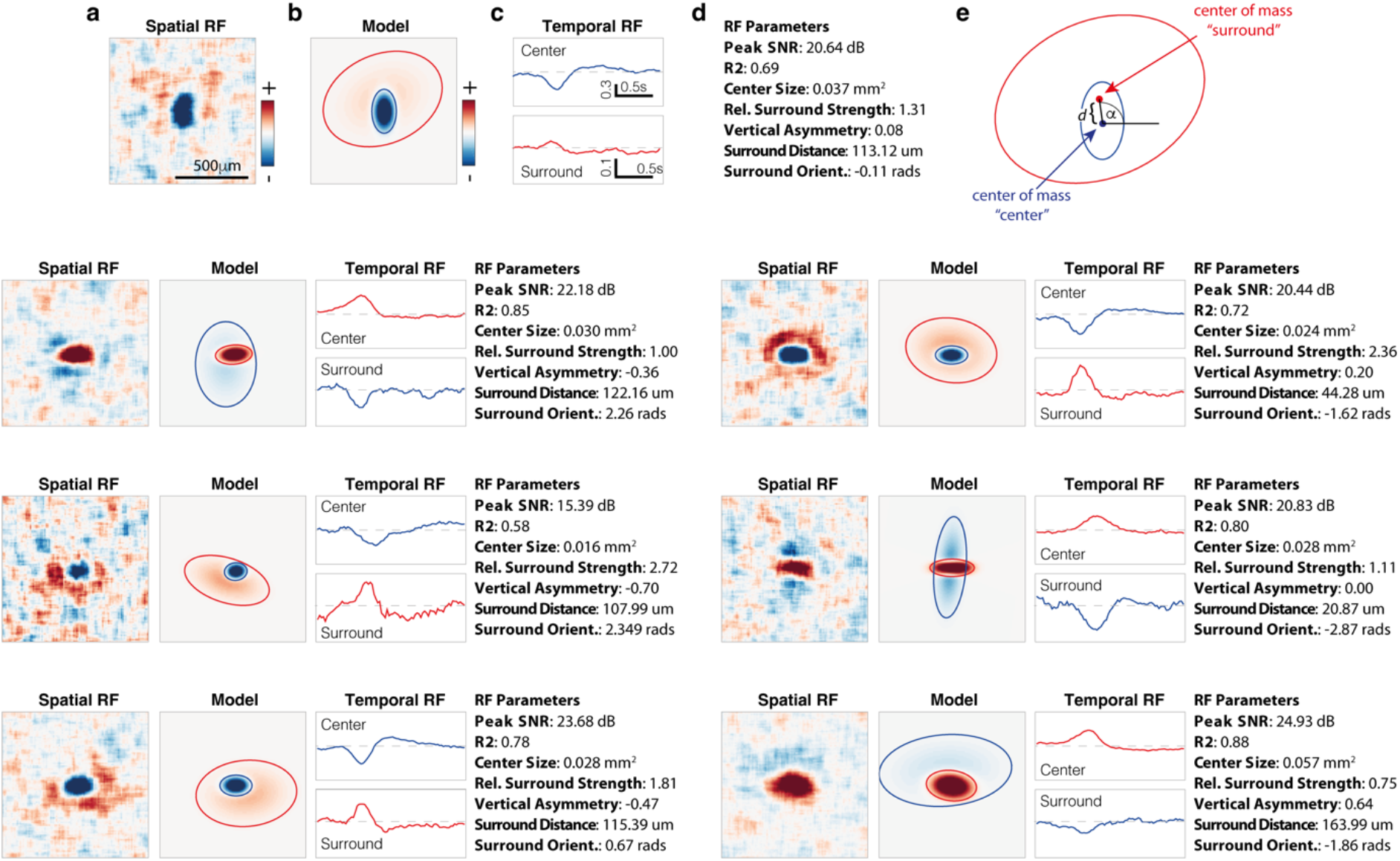
Receptive field parametrizations for representative neurons. (**a**) Spatial receptive field snapshot at the peak center strength. (**b**) Difference of Gaussians model fitted to the spatial RF. Ellipses represent 2SD of the two Gaussians. (**c**) Temporal trace of the mean value of pixels within the respective Gaussians (top: center, bottom: surround). Dashed line represents no correlation between stimulus and response. (**d**) Values of different parameters of the RF reconstruction (Peak SNR), the goodness of fit (R2), center-surround structure (Center Size, Rel. Surround Strength) and eccentricity of surround (Vertical Asymmetry, Surround Distance, Surround Orientation). All neurons are plotted with the same scale and limits (after normalization). (**e**) Schematic depicting the parametrization of the receptive field in (a), showing the surround distance (“d”) and orientation (“**α**”).

**Supplementary Fig. 6.**
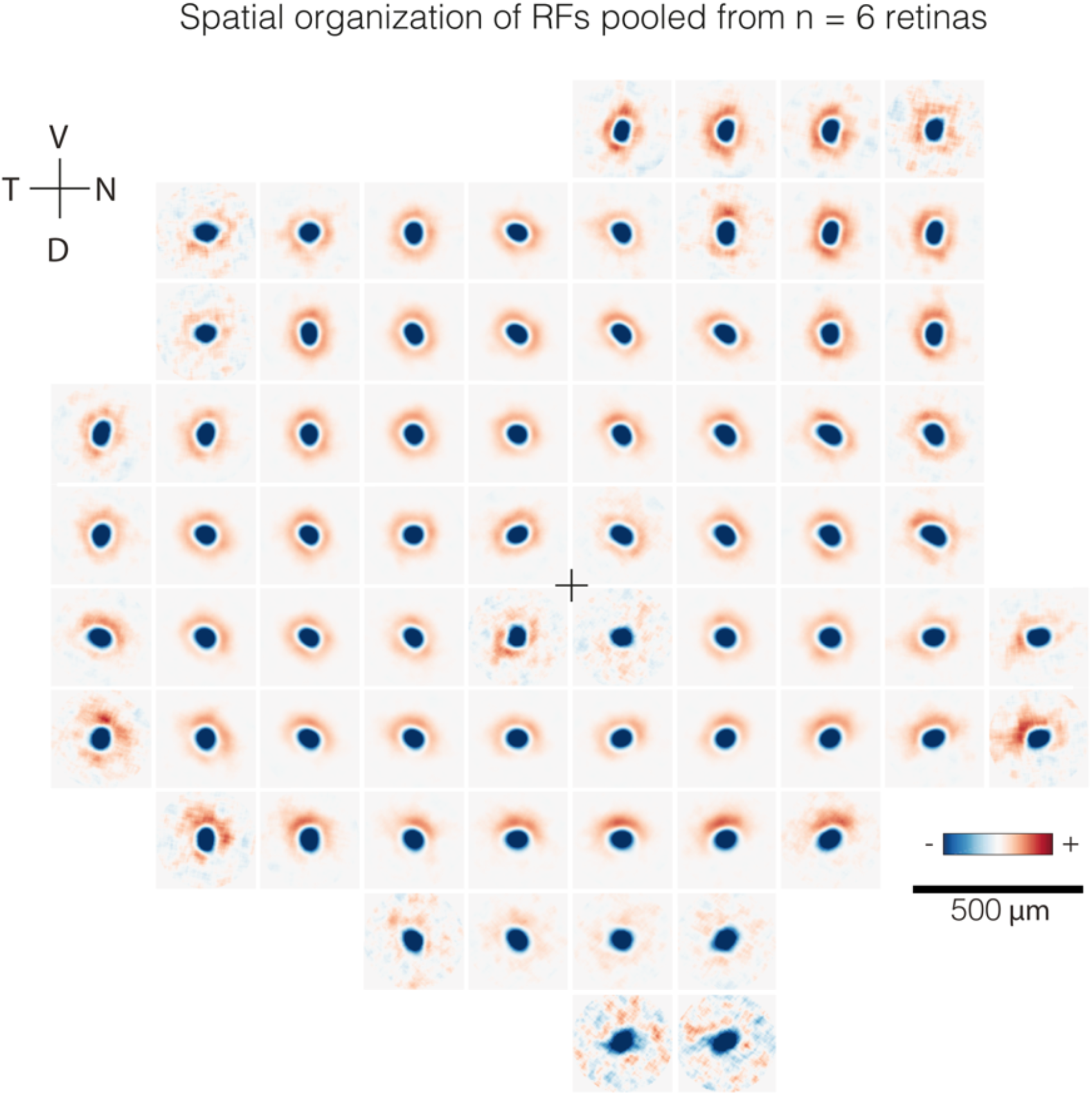
Spatial structure of average RFs across the retina. Average spatial RFs of all RGCs in square bins of size 300 μm at different positions of the retinal surface. As in Fig. 3a, but including cells from 6 retinas (n = 220 ± 200 cells per bin). Black cross: optic nerve head position.

**Supplementary Fig. 7.**
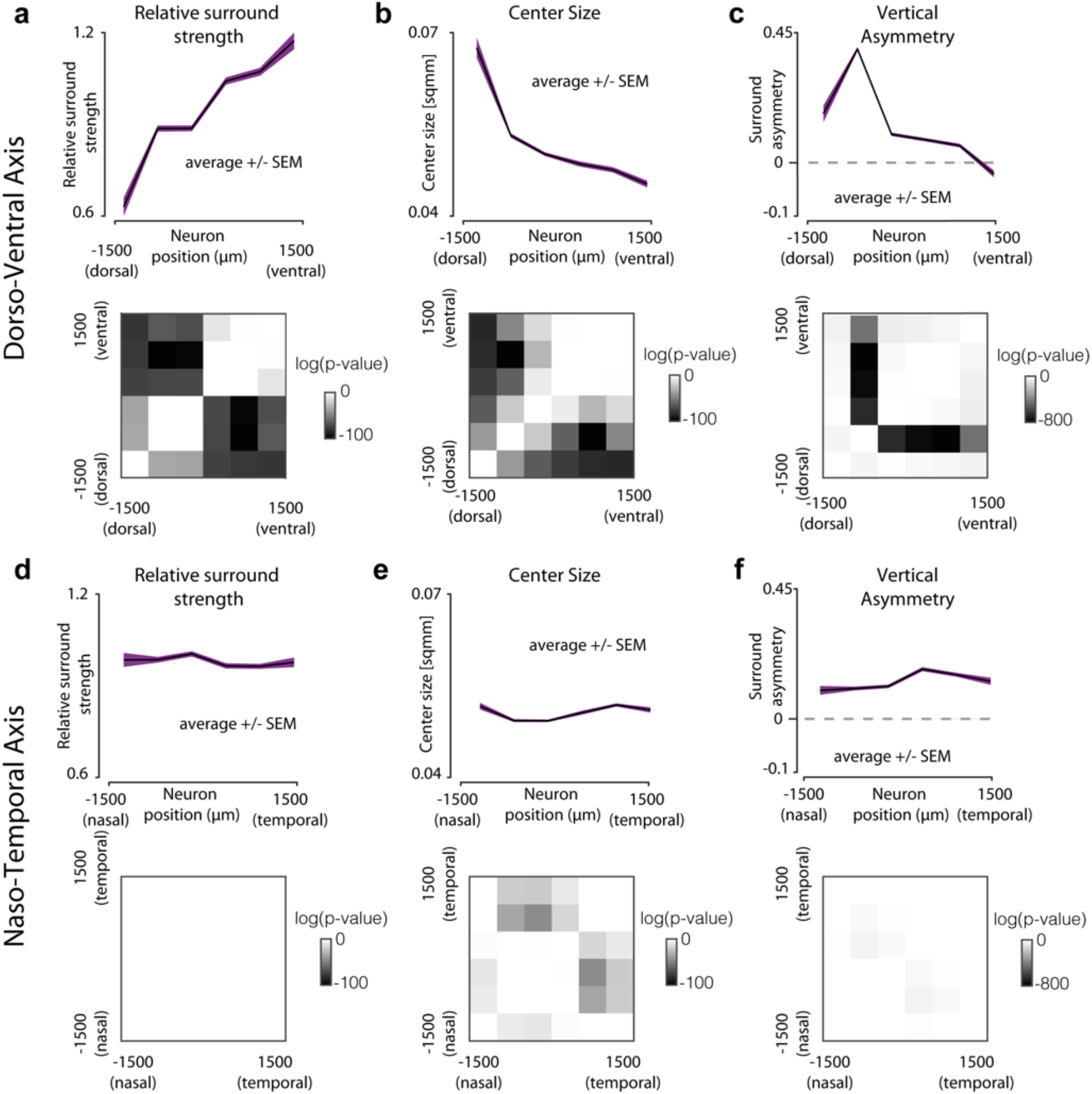
Homogeneity of receptive field architecture in the temporal-nasal axis. (**a**) Top: Mean relative surround strength at 6 different dorsoventral positions. Bottom: p-values of 2 sample Kolmogorov-Smirnov tests (with Bonferroni correction) between all pairs of bins. Darker colors represent higher significance levels that the cells in the two corresponding bins have different surround strengths. (**b**) Same as (a), but for trends and significance levels across the naso-temporal axis. (**c,d**) Same as (a,b), respectively, but for center sizes. **(e,f**) Same as (a,b), respectively, but for Vertical surround asymmetry (n = 6 retinas with S-opsin staining).

**Supplementary Fig. 8.**
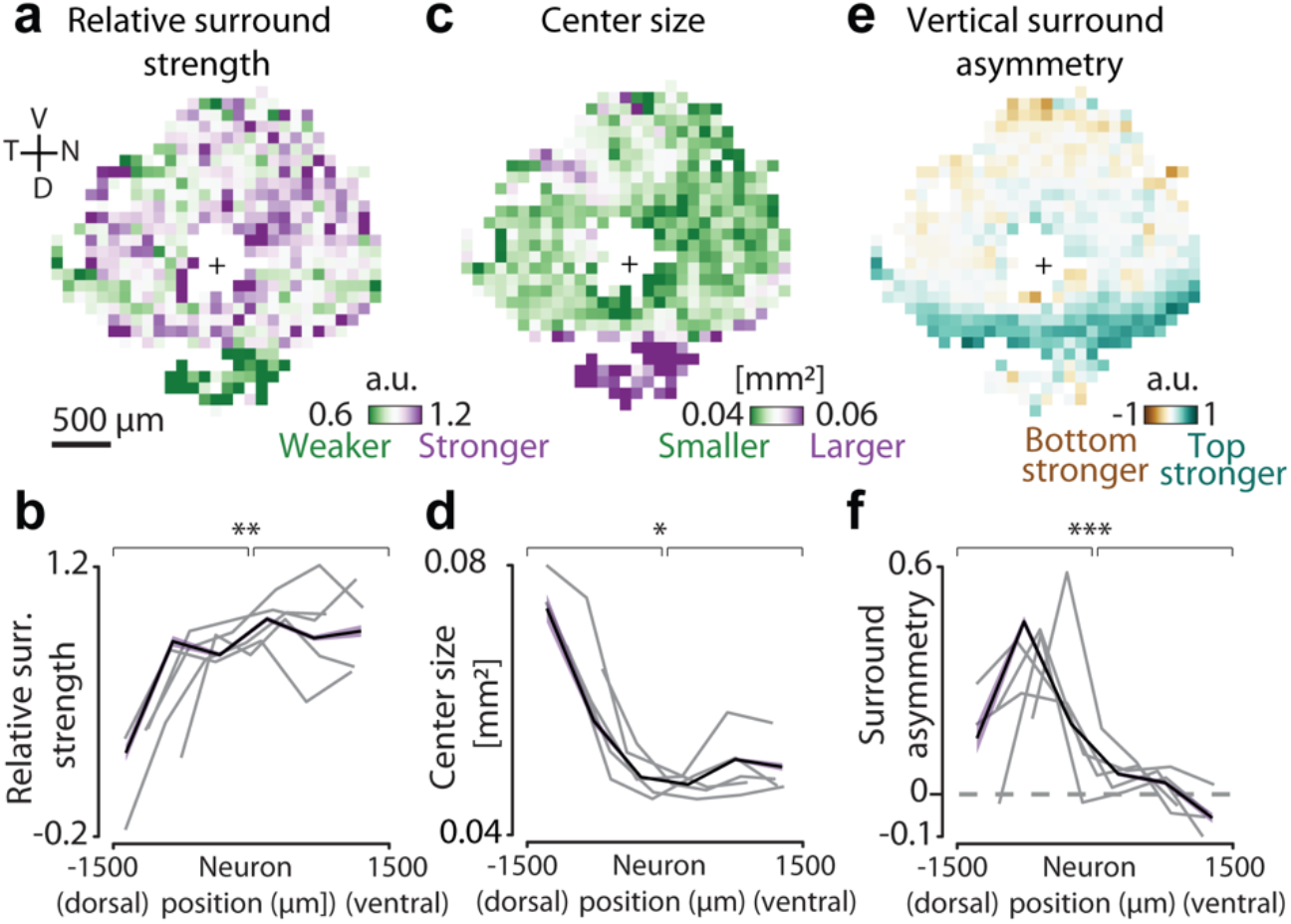
Surround strength, center size and asymmetric streak align in retinas without an S-opsin staining. (**a**) Mean relative surround strengths of RGCs within 100 μm bins, pooled from n = 5 retinas. (Same as fig 3d) (**b**) Relative surround strengths for RGCs within 6 equally spaced bins along the presumed dorsoventral axis (color: mean and SEM pooled from n = 15449 RFs, grey lines: individual retinas). (**c** & **e**) Same as (a), but for center size and vertical surround asymmetry, respectively. (**d** & **f**) same as (b), but for center size and vertical surround asymmetry, respectively. (p-values for Kolmogorov-Smirnov test: (**b**) 0.0047, (**d**): 0.0168, (**f**): 2.3766e-04;

**Supplementary Fig. 9.**
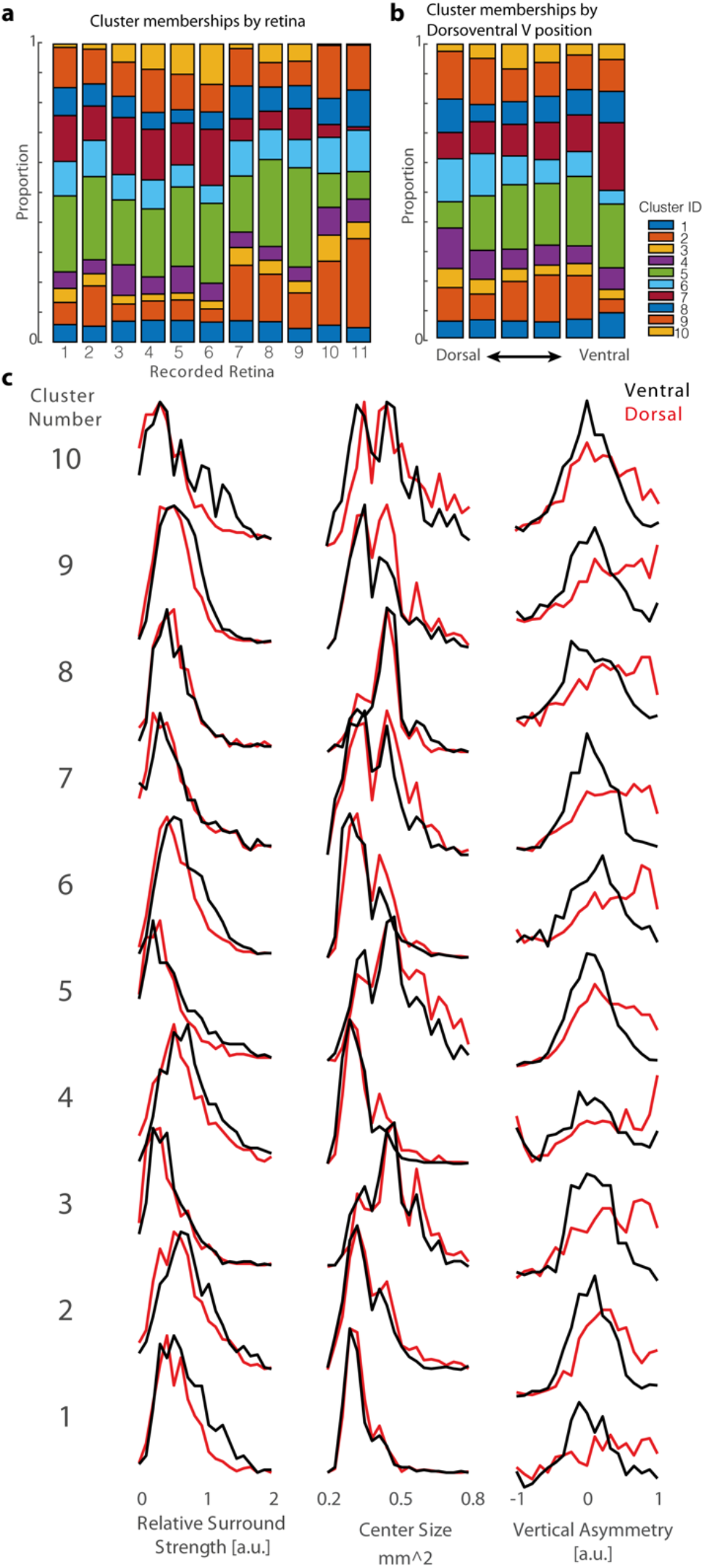
Proportions of cluster membership. (**a**) Fraction of cells from each retina that were classified into each of the 10 temporal RF clusters (from Fig. 4). (**b**) Fraction of cells across dorsoventral positions within bins of width 500 μm. (**c**) Distribution of relative surround to center strength, center sizes and vertical asymmetry for each cluster, subdivided into ventral (above the optic nerve) and dorsal (below the optic nerve). For all statistics, see Supplementary Table 1.

**Supplementary Fig. 10.**
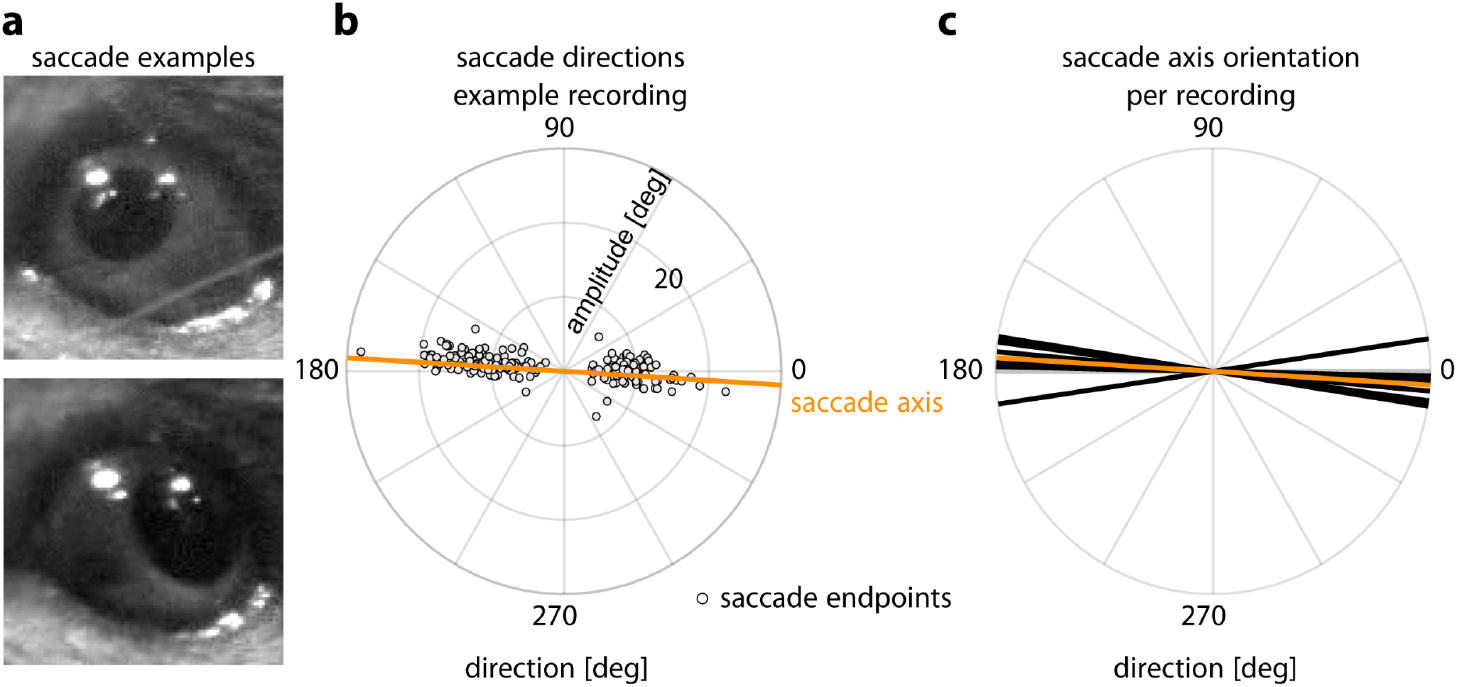
Saccades in head-fixed mice are on one axis. (**a**) Two example video frame crops of eye in two extreme horizontal positions. (**b**) Polar scatter plot of individual saccade amplitudes and directions (gray dots) in example recording and computed saccade axis (orange line, saccadic orientation tuning 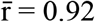). (**c**) Saccade axes of all recordings (n = 10 in 3 mice) close to the horizon (offset = 5.5 ± 2.9 °)

**Supplementary Fig. 11.**
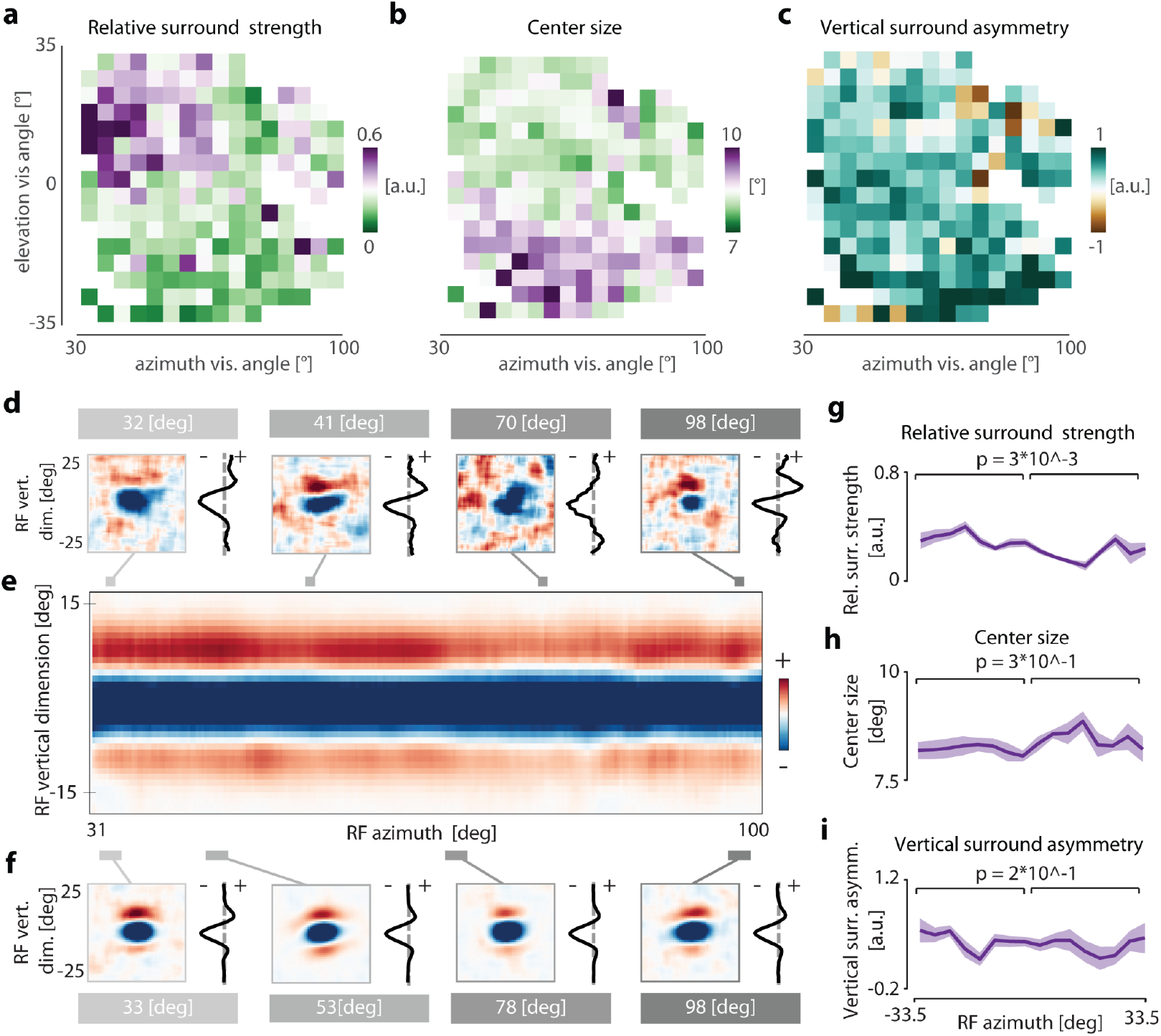
RGC bouton receptive field architecture across superior colliculus varies in elevation but not azimuth. (**a**) Relative surround strengths of mean 1-d-RFs within 4.3 ° bins. n = 45 ± 40 boutons per bin, only bins with n >= 5 boutons shown. (**b** & **c**) Same as (a), but for center size and vertical surround asymmetry, respectively. (**d**) Example RGC bouton receptive fields recorded using “shifting” white noise (left) and their respective 1-d RFs (right) at different azimuth positions (gray lines). (**e**) Average 1-d RFs, in smoothed 0.22 ° bins over azimuth. (**f**) Example average receptive fields binned at 2.9 ° visual angle (left), with their respective 1-d RFs (right) at different azimuth positions (gray bars). (**g**) Relative surround strength of 4.1 ° binned average 1-d RFs, shading indicating SEM across elevation bins (shown in (a)). Regression weights for both elevation and azimuth shown in Fig. 5g. (**h, i**) As (g), but for center size and vertical asymmetry, respectively.

**Supplementary Fig. 12.**
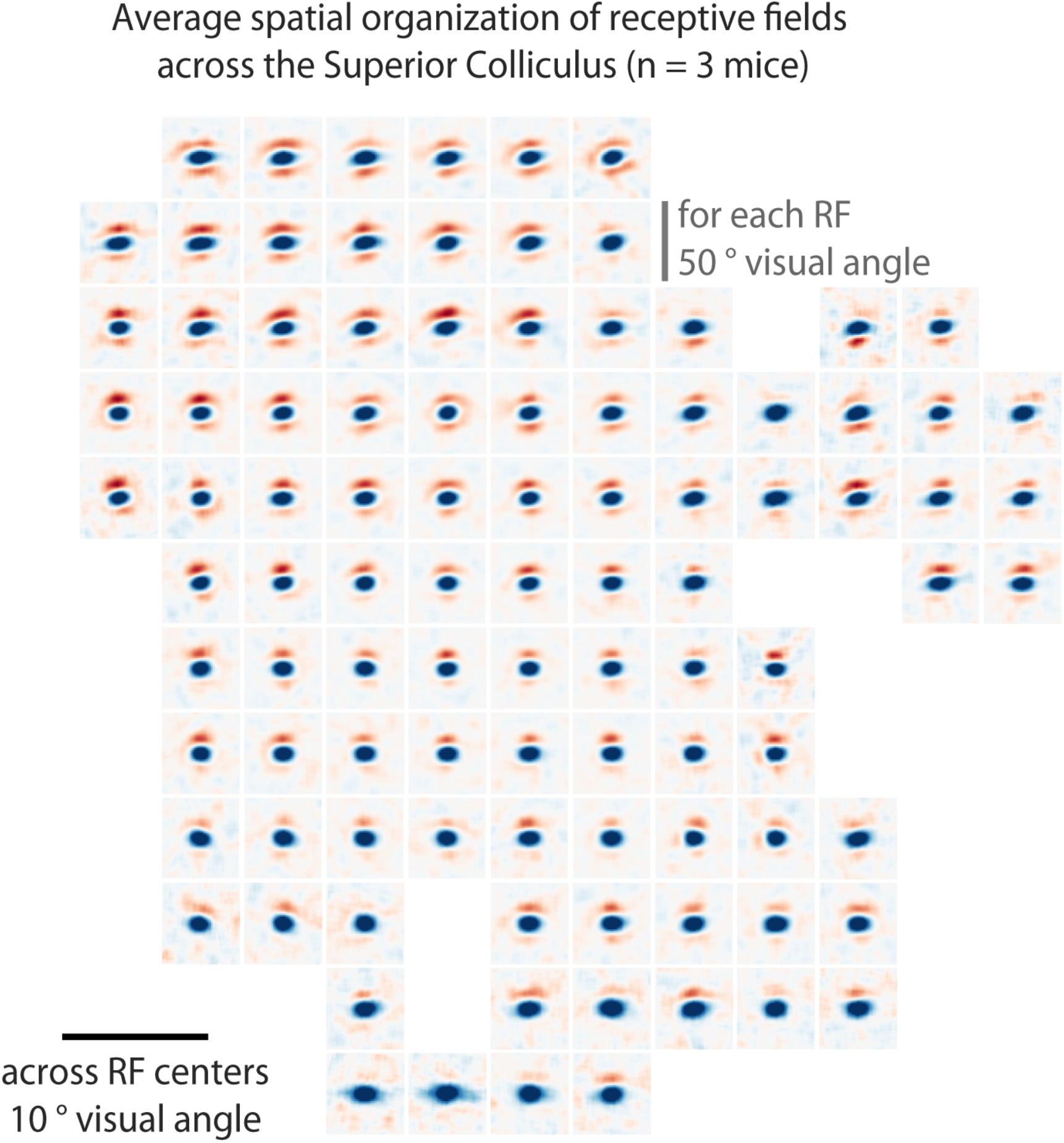
Average receptive field organization of RGC boutons across the superior colliculus. Average spatial RFs of all RGC boutons in square bins of size 5.6 ° at different positions in visual space. n = 93 ± 68 boutons per bin, only bins with n >= 20 boutons shown.

**Supplementary Table 1.**
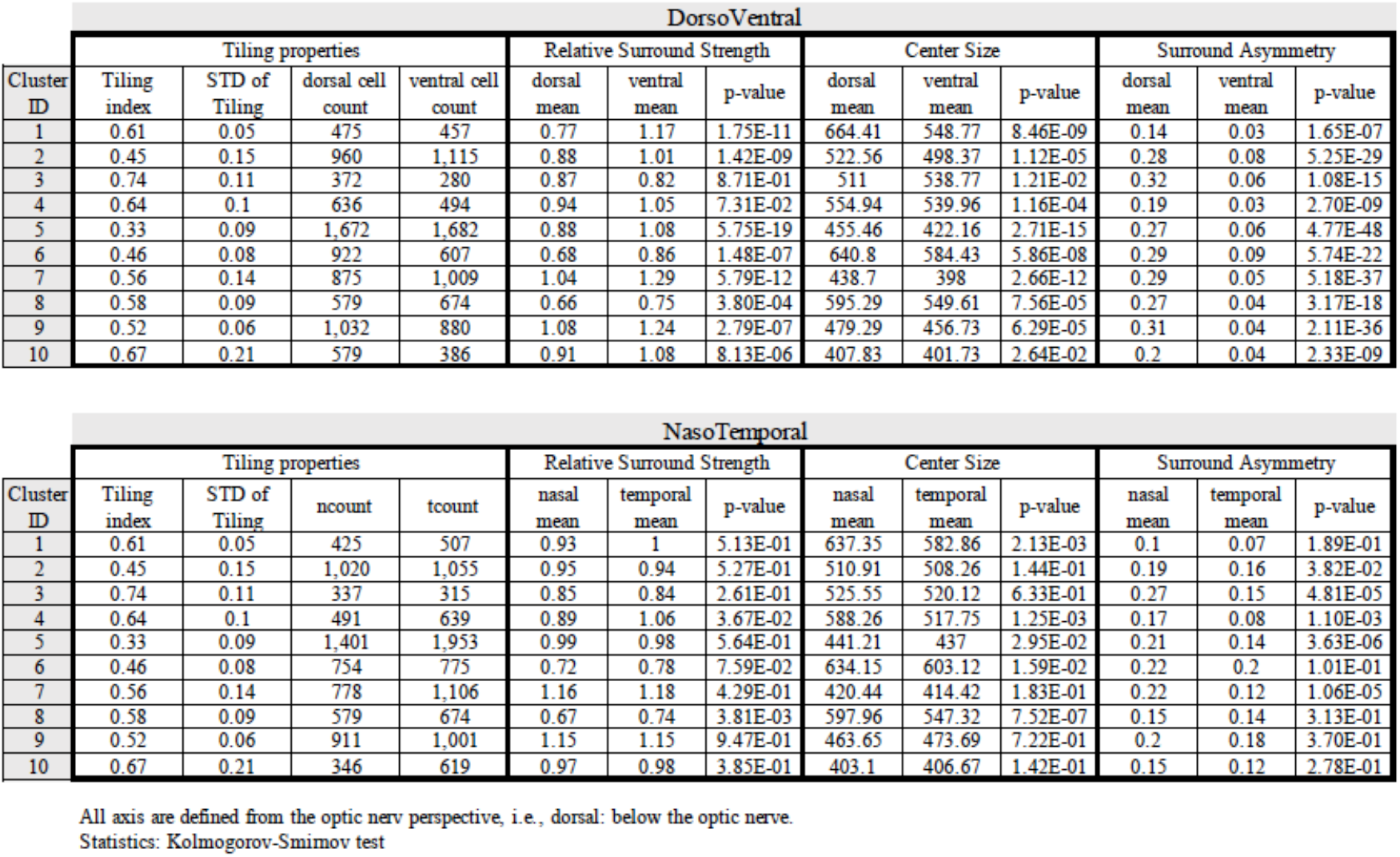
Statistics of cluster analysis for Fig. 4.

| DorsoVentral |  |  |  |  |  |  |  |  |  |  |  |  |  |
| --- | --- | --- | --- | --- | --- | --- | --- | --- | --- | --- | --- | --- | --- |
| Cluster ID | Tiling properties |  |  |  | Relative Surround Strength |  |  | Center Size |  |  | Surround Asymmetry |  |  |
|  | Tiling index | STD of Tiling | dorsal cell count | ventral cell count | dorsal mean | ventral mean | p-value | dorsal mean | ventral mean | p-value | dorsal mean | ventral mean | p-value |
| 1 | 0.61 | 0.05 | 475 | 457 | 0.77 | 1.17 | 1.75E-11 | 664.41 | 548.77 | 8.46E-09 | 0.14 | 0.03 | 1.65E-07 |
| 2 | 0.45 | 0.15 | 960 | 1,115 | 0.88 | 1.01 | 1.42E-09 | 522.56 | 498.37 | 1.12E-05 | 0.28 | 0.08 | 5.25E-29 |
| 3 | 0.74 | 0.11 | 372 | 280 | 0.87 | 0.82 | 8.71E-01 | 511 | 538.77 | 1.21E-02 | 0.32 | 0.06 | 1.08E-15 |
| 4 | 0.64 | 0.1 | 636 | 494 | 0.94 | 1.05 | 7.31E-02 | 554.94 | 539.96 | 1.16E-04 | 0.19 | 0.03 | 2.70E-09 |
| 5 | 0.33 | 0.09 | 1,672 | 1,682 | 0.88 | 1.08 | 5.75E-19 | 455.46 | 422.16 | 2.71E-15 | 0.27 | 0.06 | 4.77E-48 |
| 6 | 0.46 | 0.08 | 922 | 607 | 0.68 | 0.86 | 1.48E-07 | 640.8 | 584.43 | 5.86E-08 | 0.29 | 0.09 | 5.74E-22 |
| 7 | 0.56 | 0.14 | 875 | 1,009 | 1.04 | 1.29 | 5.79E-12 | 438.7 | 398 | 2.66E-12 | 0.29 | 0.05 | 5.18E-37 |
| 8 | 0.58 | 0.09 | 579 | 674 | 0.66 | 0.75 | 3.80E-04 | 595.29 | 549.61 | 7.56E-05 | 0.27 | 0.04 | 3.17E-18 |
| 9 | 0.52 | 0.06 | 1,032 | 880 | 1.08 | 1.24 | 2.79E-07 | 479.29 | 456.73 | 6.29E-05 | 0.31 | 0.04 | 2.11E-36 |
| 10 | 0.67 | 0.21 | 579 | 386 | 0.91 | 1.08 | 8.13E-06 | 407.83 | 401.73 | 2.64E-02 | 0.2 | 0.04 | 2.33E-09 |

| NasoTemporal |  |  |  |  |  |  |  |  |  |  |  |  |  |
| --- | --- | --- | --- | --- | --- | --- | --- | --- | --- | --- | --- | --- | --- |
| Cluster ID | Tiling properties |  |  |  | Relative Surround Strength |  |  | Center Size |  |  | Surround Asymmetry |  |  |
|  | Tiling index | STD of Tiling | ncount | tcount | nasal mean | temporal mean | p-value | nasal mean | temporal mean | p-value | nasal mean | temporal mean | p-value |
| 1 | 0.61 | 0.05 | 425 | 507 | 0.93 | 1 | 5.13E-01 | 637.35 | 582.86 | 2.13E-03 | 0.1 | 0.07 | 1.89E-01 |
| 2 | 0.45 | 0.15 | 1,020 | 1,055 | 0.95 | 0.94 | 5.27E-01 | 510.91 | 508.26 | 1.44E-01 | 0.19 | 0.16 | 3.82E-02 |
| 3 | 0.74 | 0.11 | 337 | 315 | 0.85 | 0.84 | 2.61E-01 | 525.55 | 520.12 | 6.33E-01 | 0.27 | 0.15 | 4.81E-05 |
| 4 | 0.64 | 0.1 | 491 | 639 | 0.89 | 1.06 | 3.67E-02 | 588.26 | 517.75 | 1.25E-03 | 0.17 | 0.08 | 1.10E-03 |
| 5 | 0.33 | 0.09 | 1,401 | 1,953 | 0.99 | 0.98 | 5.64E-01 | 441.21 | 437 | 2.95E-02 | 0.21 | 0.14 | 3.63E-06 |
| 6 | 0.46 | 0.08 | 754 | 775 | 0.72 | 0.78 | 7.59E-02 | 634.15 | 603.12 | 1.59E-02 | 0.22 | 0.2 | 1.01E-01 |
| 7 | 0.56 | 0.14 | 778 | 1,106 | 1.16 | 1.18 | 4.29E-01 | 420.44 | 414.42 | 1.83E-01 | 0.22 | 0.12 | 1.06E-05 |
| 8 | 0.58 | 0.09 | 579 | 674 | 0.67 | 0.74 | 3.81E-03 | 597.96 | 547.32 | 7.52E-07 | 0.15 | 0.14 | 3.13E-01 |
| 9 | 0.52 | 0.06 | 911 | 1,001 | 1.15 | 1.15 | 9.47E-01 | 463.65 | 473.69 | 7.22E-01 | 0.2 | 0.18 | 3.70E-01 |
| 10 | 0.67 | 0.21 | 346 | 619 | 0.97 | 0.98 | 3.85E-01 | 403.1 | 406.67 | 1.42E-01 | 0.15 | 0.12 | 2.78E-01 |

All axis are defined from the optic nerv perspective, i.e., dorsal: below the optic nerve. Statistics: Kolmogorov-Smirnov test

## Notes

### Competing Interest Statement

The authors have declared no competing interest.

### Summary of Updates

We have added substantial new data and analysis from both the experimental and theoretical sides. Most importantly, we have added a new figure to the manuscript showing that our main experimental result, the efficient changes in receptive field structures, is preserved in retinal ganglion cell terminals of awake-behaving animals that have an intact visual system.

